# Cretaceous crocodyliform reconciles conflicting evidence on the Mesozoic paleogeography of Europe during the Gondwana-Laurasia split

**DOI:** 10.1101/2025.08.25.671775

**Authors:** Máté Szegszárdi, Attila Ősi, Márton Rabi

## Abstract

Inferred biotic exchanges between Europe and Africa during the Cretaceous have been used to support the hypothesis that the complete separation of Europe from Gondwana postdated the split of the remaining Laurasian landmasses from Gondwana. Under this framework, Europe, conceptualized as part of a proposed ‘Eurogondwana’, is thought to have maintained biogeographic connections with Africa well into the Cretaceous. A key piece of evidence underpinning this hypothesis has been the crocodyliform *Doratodon carcharidens* from the Late Cretaceous of Europe, previously interpreted as closely related to Gondwanan taxa. However, the material attributed to this species is fragmentary, and its skeletal anatomy—critical for phylogenetic inference—remains poorly understood. Here we describe a new partial skull of *Doratodon carcharidens* which represents some of the most complete material of ‘Gondwanan’ taxa in the European Cretaceous. Our updated phylogenetic analysis unexpectedly and robustly places *D. carcharidens* among the Laurasian clade Paralligatoridae and reveals that morphological similarities to Gondwanan ziphosuchians, such as ziphodont dentition, are the result of ecomorphological convergence rather than shared ancestry. By reinterpreting the systematic position of further taxa representing biogeographic enigma, such as *Ogresuchus furatus*, our phylogeny implies a major role of the primary breakup of Pangea into Laurasia and Gondwana for crocodyliform divergence. A critical reassessment of the purported evidence for ‘Gondwanan’ fish and tetrapod immigrants in Europe reveals that it is largely based on highly fragmentary and sporadic specimens, as well as weakly supported phylogenetic hypotheses. Given the sparse and uneven Jurassic and Cretaceous fossil record in both Europe and Africa, it remains plausible that taxa previously interpreted as Gondwanan dispersers instead represent vicariant relicts. Our results conflict with recent paleobiogeographic scenarios, highlight the absence of compelling evidence for the Eurogondwana hypothesis and instead support a primary Gondwana-Laurasia split.

## Introduction

The Mesozoic primary split of Pangea into Laurasia and Gondwana has been a longstanding paleogeographic paradigm supported by geological, fossil, and molecular evidence [1,2]. However, this has been increasingly challenged by more recent studies on European and North African Cretaceous continental faunas, which are of critical interest for paleogeography since they constitute independent evidence for land connections and proximity between the two continents [3,4,5,6]. Particularly, the presence of tetrapod clades with purported Gondwanan affinities in the European record have been difficult to reconcile with a Laurasia-Gondwana dichotomy and has been explained by unidirectional dispersal events, cosmopolitanism [3,7,8,9], or more recently, by the retention of faunal links between Europe and Gondwana well after the initial breakup of Pangea [5,10]. The latter concept postulates a primary dichotomy between Asiaamerica and a combined Gondwana-Europe supercontinent, ‘Eurogondwana’, in the earliest Cretaceous mostly based on archosaur (dinosaurs and crocodyliforms) and to lesser extent other tetrapod faunal evidence. According to this model, Laurasia, including Europe, would not have formed before the Hauterivian and was only isolated till the end of the Cretaceous when extensive biotic interchange between Europe and Gondwana “renewed” via the Apulian microcontinent [10]. The main appeal of the ‘Eurogondwana’ model is that it provides an explanation for the absence of Laurasian taxa in the Gondwanan Cretaceous up until the Campanian-Maastrichtian, otherwise difficult to interpret with land bridges normally promoting bidirectional dispersals. By integrating additional tetrapod evidence, subsequent work attributed the absence of Laurasian taxa in Africa to sampling bias and revealed that at face value, the fossil record implies persistent input of ‘Gondwanan’ clades into European faunas throughout the Cretaceous [5]. Recently, new fossil finds and updated taxonomies supported the presence of Laurasian taxa in Africa and thereby seemingly filling part of the gap in the Gondwanan record [11,12,13,14]. To date, the presence of ‘Gondwanan’ tetrapods in the European Cretaceous is broadly accepted but uneven sampling makes it difficult to determine whether the biotic exchange was persistent or temporary.

Nevertheless, the ‘Eurogondwana’ hypothesis remains in strong contrast with plate tectonic models reconstructing a primary split of Laurasia and Gondwana driven by the opening of the Central Atlantic and the Alpine Tethys oceans with Africa commencing its drift towards Europe only after the subsequent opening of the southern Atlantic Ocean [15,16]. Moreover, the Cretaceous fauna of Europe was dominated by Laurasian clades and yet evidence for these taxa in Africa remains disproportionately fewer. The presence of ‘Gondwanan’ tetrapods in the European Cretaceous and the continent’s history during the primary split of Pangea is therefore far from resolved.

One of the purported ‘Gondwanan’ taxa of key importance to the Eurogondwana model has been the crocodyliform *Doratodon carcharidens* [17], known by fragmentary cranial remains from the lower Campanian Gosau Group of Austria and the Santonian Csehbánya Formation of Hungary [5]. *Doratodon carcharidens* is characterized by ziphodont teeth, a deep, laterally compressed snout, closed mandibular fenestra, and the presence of an antorbital fenestra [5,18,19]. Previous phylogenies placed this taxon closely related to South American members of the Gondwanan ziphosuchian clade Sebecosuchia [5,20], a group primarily known for their ziphodont dentition and oreinirostral snout [21,22]. *D. carcharidens* represents one of the few taxa used in support for the Gondwanan affinity of the European Cretaceous fauna based on phylogenetic evidence [5,9,13,23]. The osteology of *D. carcharidens* nevertheless remains poorly known and has been limited to jaw remains and teeth.

We here report an incomplete skull and additional isolated cranial elements of *Doratodon carcharidens* from the Santonian Csehbánya Formation of Hungary [24,25] and by rigorously testing its phylogenetic relationships based on these specimens using multiple comprehensive taxon-character datasets, we demonstrate that the shared traits with Sebecosuchians is due to ecomorphological convergence and this taxon is more closely related to Laurasian Neosuchian crocodyliforms. The unexpected systematic reinterpretation of *D. carcharidens* highlights the need for a critical review of the evidence used to infer vertebrate dispersals between Europe and Gondwana during the Cretaceous. We find that both the preservation and temporal sampling of the relevant taxa is highly incomplete, which questions the reliability of the phylogenies underpinning the Eurogondwana model.

## Results

### Systematic paleontology

Crocodyliformes Hay, 1930

Mesoeucrocodylia Whetstone & Whybrow, 1983

Neosuchia Benton & Clark, 1988 [26]

Paralligatoridae Konzhukova, 1954 [27]

*Doratodon* Seeley, 1881 [18]

*Doratodon carcharidens* Bunzel, 1871 [17]

#### Holotype

IPUW 2349/57, almost complete mandible.

#### Type locality and horizon

Konstantin mining tunnel, Felbering Mine, Muthmannsdorf, Wiener Neustadt-Land district, Niederösterreich (Lower Austria), Austria, Grünbach Formation, Gosau Group, lower Campanian.

#### Referred specimens

Material from the Austrian type locality [5,17,18,19]: maxilla (IPUW2349/5), parietal (UWPI 2349/54); material from the Santonian Csehbánya Formation, Iharkút, Hungary (Rabi & Sebők [5], this study): partial skull (MTM PAL 2024.159.1, Figs. 1-3.) including the posterior portion of the skull, paired nasals, pterygoid, right maxilla and left ectopterygoid; isolated quadrate (MTM PAL 2013.67.1.), isolated pterygoid (MTM PAL 2013.64.1.) (Fig. 4.), right premaxilla (MTM PAL 2014.122.1) with in situ teeth, fragmentary left maxilla (MTM PAL 2013.65.1), fragmentary left dentary (MTM V2010.237.1), fragmentary right dentary (MTM PAL 2013.66.1).

**Fig. 1.**
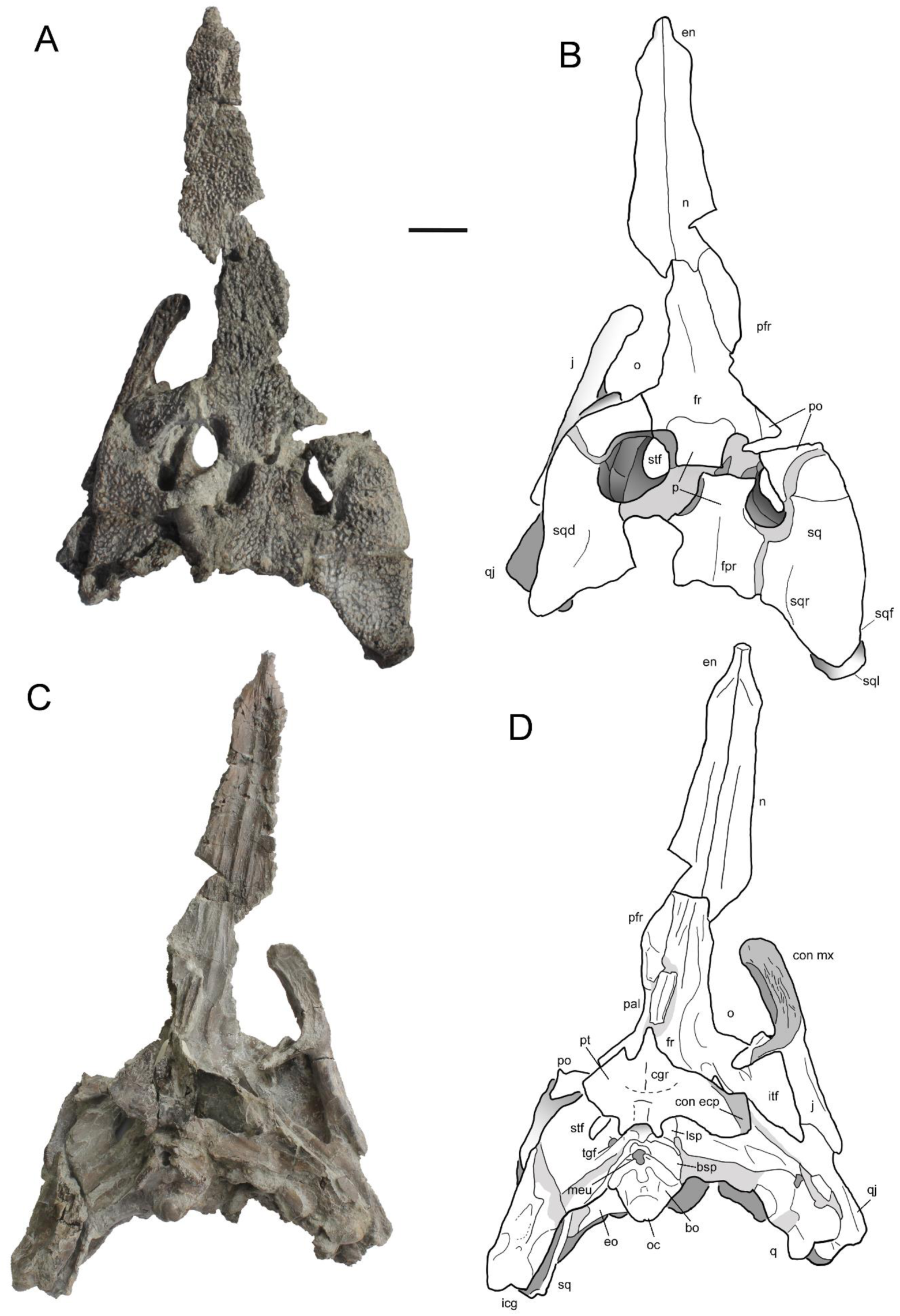
**Partial** skull (MTM PAL 2024.159.1) of ***Doratodon carcharidens* from the Santonian Csehbánya Formation of Iharkút, Hungary.** (**A**, **B**) dorsal and (**C**, **D**) ventral view. Further disarticulated elements of this specimen are figured in Figs. 3-4. Scale bar equals 2 cm. **Anatomical abbreviations:** bo, basioccipital; bsp, basisphenoid; cgr, choanal groove; ch, choana; chs, choanal septum; con ecp, contact with ectopterygoid; en, external nares; eo, exoccipital; fa, foramen aereum; fr, frontal; fpr, frontoparietal ridge; itf, infratemporal fenestra; j, jugal; lc, lateral condyle; lsp, laterosphenoid; mc, medial condyle; meu, median Eustachian foramen; mp, maxillary palate; n, nasal; o, orbit; oa, otic aperture; p, parietal; pal, palatine; pfr, prefrontal; pmpr, posteromedial process; po, postorbital; pt, pterygoid; ptw, pterygoid wing; q, quadrate; qj, quadratojugal; sq, squamosal; sqd, squamosal depression; sql, squamosal lobe; sqr, squamosal ridge; so, supraoccipital, sof, suborbital fenestra; stf, supratemporal fenestra; tgf, trigeminal foramen.

**Fig. 2.**
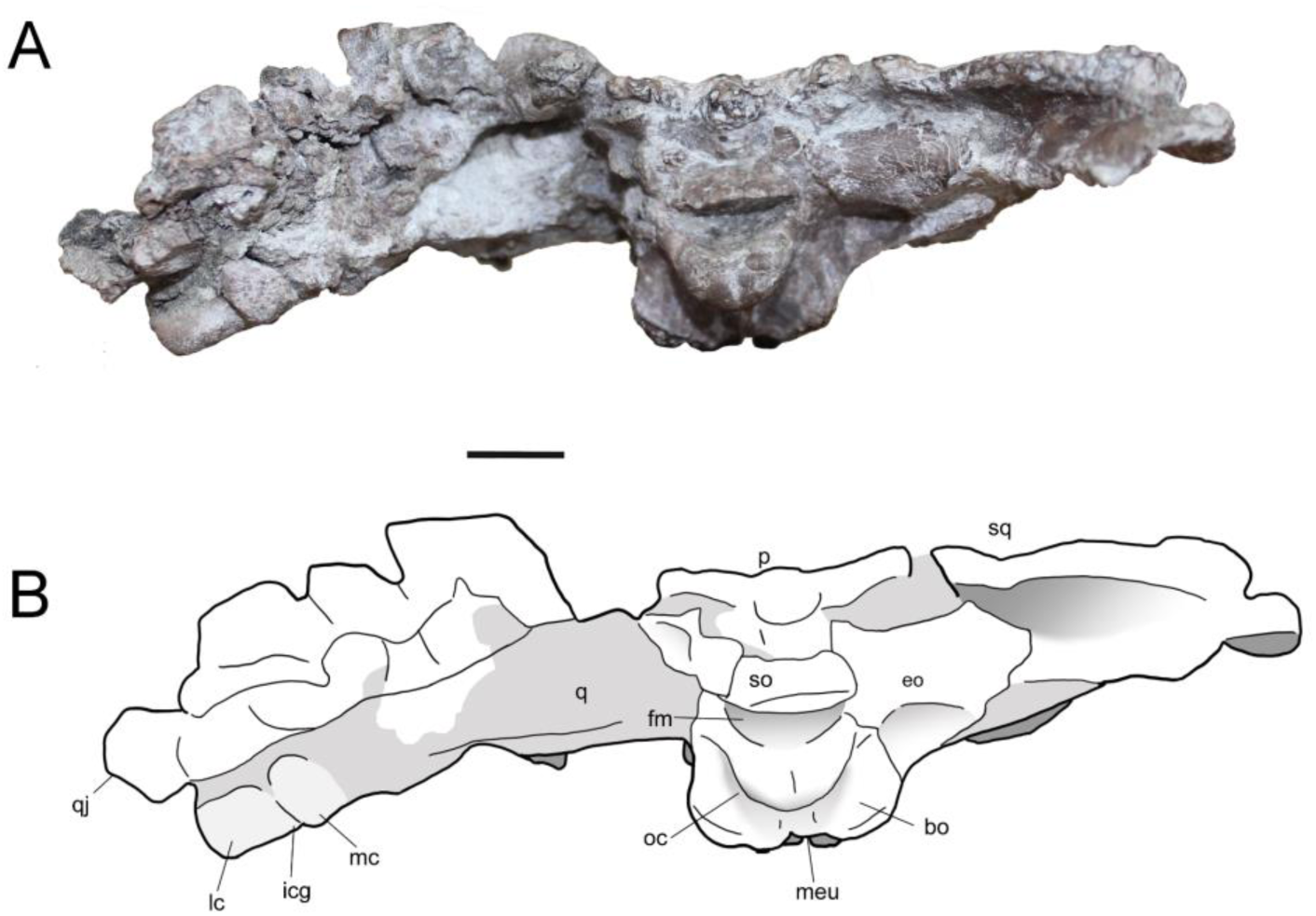
**Partial skull (**MTM PAL 2024.159.1) of the *Doratodon carcharidens* skull from the Santonian Csehbánya Formation of Iharkút, Hungary. Photograph (**A**) and interpretive line drawing (**B**) in occipital view. Scale bar equals 1 cm. **Anatomical abbreviations:** bo, basioccipital; bsp, basisphenoid; eo, exoccipital; fm, foramen magnum; icg, intercondylar groove; lc, lateral condyle of condylus mandibularis of quadrate; mc, medial condyle of condylus mandibularis of quadrate; meu, median Eustachian foramen; oc, occipital condyle; p, parietal; po, postorbital; pt, pterygoid; ptw, pterygoid wing; q, quadrate; qj, quadratojugal; s, squamosal; sql, squamosal lobe; so, supraoccipital; tgf, trigeminal foramen.

**Fig. 3.**
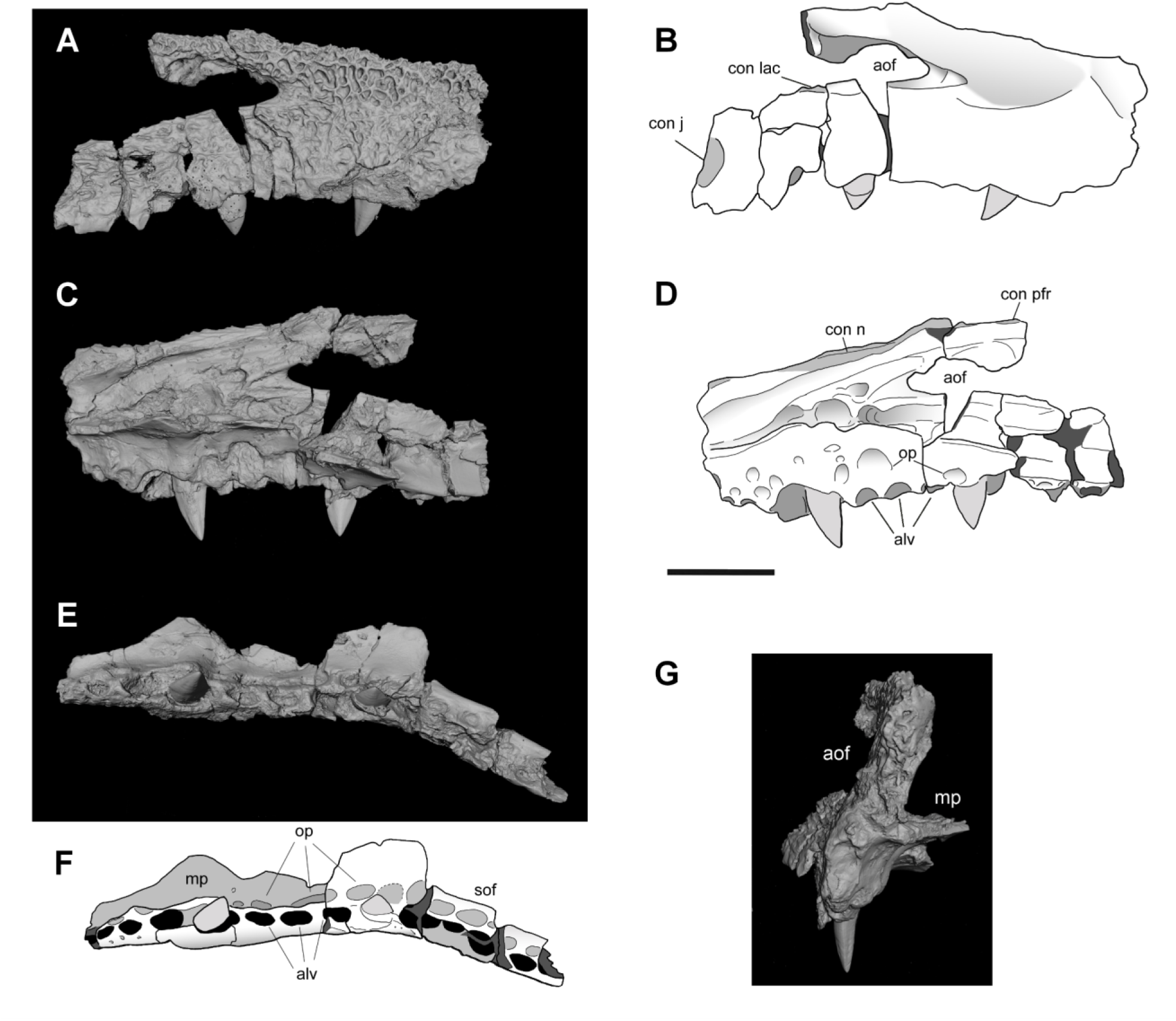
**Disarticulated maxilla of the** MTM PAL 2024.159.1 *Doratodon carcharidens* skull from the Santonian Csehbánya Formation of Iharkút, Hungary.(**A**, **B**), medial (**C**, **D**), ventral (**E**, **F**) and anterior (**G**) view. Scale bar equals 2 cm. **Anatomical abbreviations:** aof, antorbital fenestra; con j, contact with jugal; con l, contact with lacrimal; con n, contact with nasal; con pfr, contact with prefrontal; mp, maxillary palate.

**Fig. 4.**
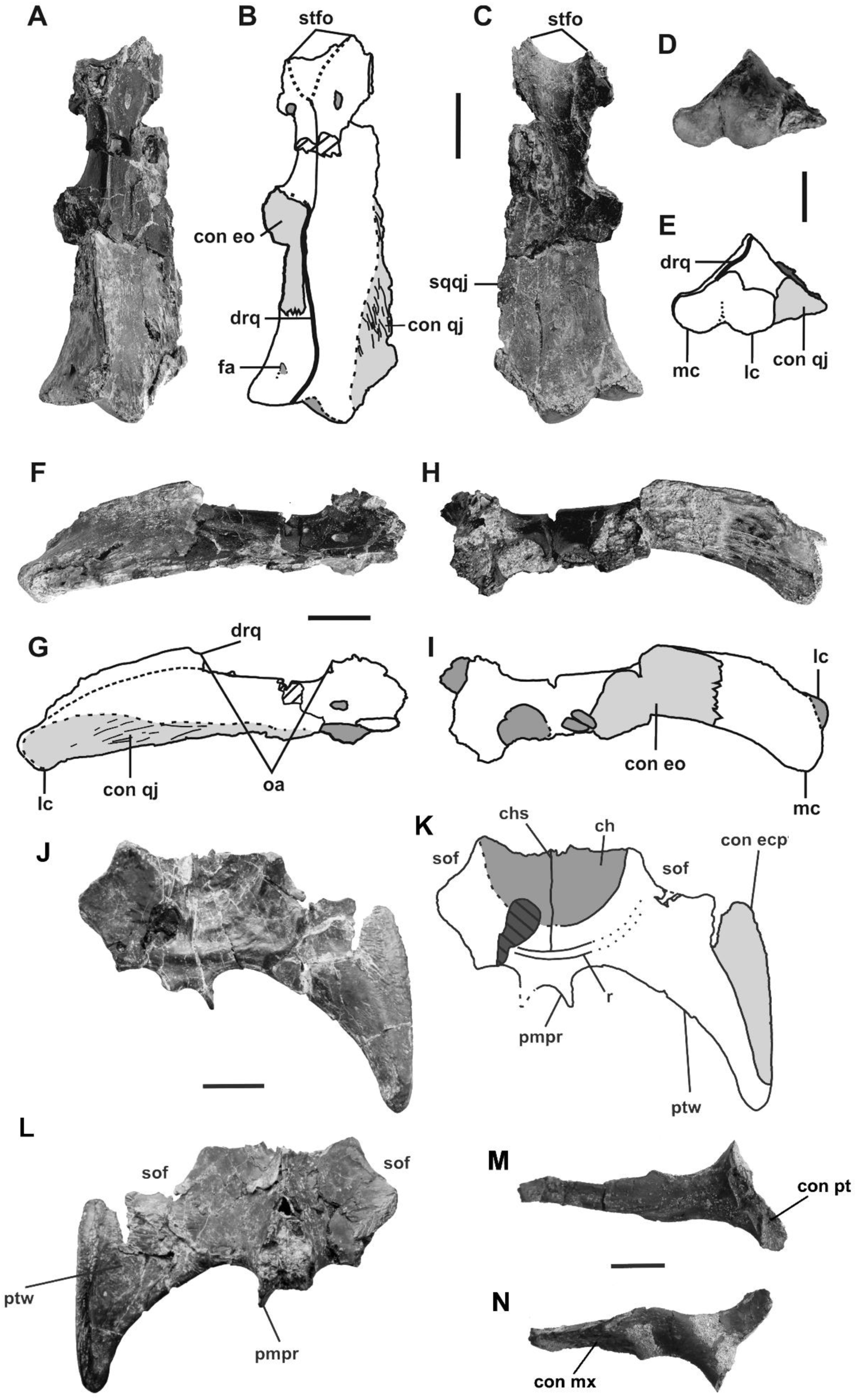
*Doratodon carcharidens* cranial remains from the Santonian Csehbánya Formation of Iharkút, Hungary. Photographs and interpretive drawings of MTM PAL 2013.67.1. quadrate in dorsal (**A**, **B**), ventral (**C**), posterior (**D**, **E**), lateral (**F**, **G**), and medial view (**H**, **I**); **J**-**L**: Photographs and interpretive drawing of MTM PAL 2013.64.1. pterygoid in ventral (**J**, **K**,) and dorsal view (**L**); **M**-**N**: ectopterygoid of l MTM PAL 2024.159.1 skull (Figs. 1-3) in medial (**M**) and lateral (**N**) view. Light gray denotes sutural surfaces, while dark gray indicates foramina and depressions. All scale bars equal 1 cm. Photographs and illustrations **A**-**L** by Nóra Sebők. **Anatomical abbreviations:** cgr, choanal groove; ch, choana; chs, choanal septum; con ecp, contact with ectopterygoid; con pt, contact with pterygoid; con qj, contact with quadratojugal; drq, dorsal ridge of quadrate; fa, foramen aereum; lc, lateral condyle; mc, medial condyle; oa, otic aperture; pmpr, posteromedial process of pterygoid; ptw, pterygoid wing; sof, suborbital fenestra.

#### Revised diagnosis

Small-sized paralligatorid with a narrow rostrum, a deep, subvertical maxilla with straight alveolar edges, and an antorbital fenestra. The maxilla and dentary bear circular pits near the alveolar surface, medial to the tooth row in the maxilla and laterally in the dentary. Distinguished from other paralligatorids on the basis of a present but reduced antorbital fenestra, a longitudinal lateral depression on the anterior region of the surangular and the posterior region of the dentary, the external maxillary surface forming a single laterally facing plane, a distinct posterior and medial face on the quadrate in posterior view, the latter of which bears the foramen aereum, a straight ventral maxillary edge, and a cranial table about as wide as the ventral portion of the skull, all of which are recovered as autapomorphies in the phylogenetic analyses. It differs from all other paralligatorids except *Wannchampsus kirpachi* in bearing denticulate carinae on the teeth. *Doratodon carcharidens* differs from *D. ibericus* in a mandibular tooth row that is more homodont in size, a less abrupt slope in the symphyseal area, and a single diastema between the 2nd and 3rd alveoli, as in Company et al. [28].

#### Remarks

The skull MTM PAL 2024.159.1 (Figs. 1-3) is referred to *Doratodon carcharidens*, a taxon previously reported from the Iharkút locality, based on the subvertical lateral surface of the maxilla, presence of an antorbital fenestra, a row of prominent circular to anteroposteriorly oval occlusal pits medial to the maxillary tooth row, and labiolingually compressed, ziphodont teeth [5]. The right maxilla of MTM PAL 2024.159.1 (Fig. 3) was found isolated from the rest of the skull within a distance of 150-200 cm and its contact surface and ornamentation matches with that of the right nasal perfectly. The isolated quadrate (MTM PAL 2013.67.1.) and pterygoid (MTM PAL 2013.64.1.) are referred to *D. carcharidens* based on their consistent morphology with the equivalent elements of MTM PAL 2024.159.1. The overlapping specimens of the Iharkút and type-locality material show no apparent differences except for the number of symphyseal occlusal pits in the dentary (two instead of a single pair in the holotype) [5] and we therefore consider them conspecific.

#### Description

The paired nasals are long and narrow, possessing a downturned anterior process separating the anteriorly directed external nares. The right prefrontal borders the nasal posterolaterally and has a posteriorly deflected, gracile prefrontal bar with a concave anteroventral surface. The prefrontal bar is thick towards the prefrontal, tapers ventromedially, and becomes laminar near the palatines. The maxilla is characterized by a straight ventral edge and an outer surface formed by a single, laterally facing plane. Out of the 14 preserved maxillary alveoli, the 5th and 9th teeth are in place, although their exact position in the tooth row is unknown. The teeth are labiolingually compressed and ziphodont with slightly posteriorly curved apex, with the preserved 5th tooth being larger, more curved, and more slender than the 9th. Isolated *D. carcharidens* teeth previously reported from the same site are described in Rabi & Sebők [5]. The maxilla has a row of circular to anteroposteriorly oval occlusal pits medial to the tooth row, approximately the same size as the alveoli except for the one posteromedial to the preserved 5th and anteromedial to the 6th tooth. Based on the external nares and the maxilla, the snout was oreinirostral and anteriorly tapering. An anteroposteriorly elongated, oval antorbital fenestra is present and the maxillary palate borders the suborbital fenestra anteriorly.

The skull table has a flat, trapezoid dorsal surface laterally extending to cover most of the quadrate and quadratojugal, decorated with fine pitting. The unpaired frontal is anteroposteriorly elongated, lacks a fossa at the anteromedial margin of the supratemporal fenestra, and bears a medial longitudinal ridge, extending and terminating in a posterior protrusion on the parietal. The squamosal-parietal suture, although broken, is marked by a posterior protrusion on each side. The squamosals are subtriangular with a slightly concave posterior edge, and bear lobe-like, unsculpted ventrolateral processes that extend posteriorly. Although slightly distorted by dorsoventral compression, a flare, a ridge and a depression (see Turner [29], Fig. 4) are present on the dorsal surface and lateral edge of the squamosal.

The jugal is elongated and narrow, widening only slightly anterior to the postorbital bar. Near the maxillary suture, the lateral surface of the jugal bears a distinctive ornamentation consisting of elongated oval pits bordered by sharp edges. The concave ventral half of the lateral surface is separated from the dorsal half by a low ridge. The quadratojugal contacts the jugal through its lateral process and the quadrate along the medial margin of its medial process. A quadratojugal spine is absent. Attachment sites for the M. adductor mandibulae externus superficialis are visible near the suture with the quadrate on the ventral surface of the quadratojugal. The quadrate contacts the quadratojugal laterally, the pterygoid anteromedially, and also borders the supratemporal fossa and trigeminal fenestra. Its posteroventral end bears two distinct condyles separated by a V-shaped intercondylar groove. Based on specimen MTM PAL 2013.67.1., the quadrate was subtriangular in posterior view prior to the dorsoventral compression, with a medial and a lateral face separated by a ridge.

The pterygoid flanges project ventrolaterally, although only the left flange is preserved in its entirety, bearing a portion of the dorsoventrally expanded lateral head. Based on an isolated pterygoid of a different individual (MTM PAL 2013.64.1.), the head expanded posteriorly further than the main body of the quadrate, with a posteriorly tapering surface. The pterygoid does not fully enclose the choana; MTM PAL 2013.64.1. bears a partially septated choanal groove and an elevated posterior choanal border. The ectopterygoid recovered in association with MTM PAL 2024.159.1 has a narrow and gracile ectopterygoid bar, with a medial articular surface about twice as wide, and an elongated lateral articular surface. Its articulation to the jugal is unclear due to the absence of the right jugal in MTM PAL 2024.159.1, although the general shape of the suture with the pterygoid closely resembles the ectopterygoid suture of the MTM PAL 2013.64.1. specimen (see Fig. 4). Only a posterior fragment of the palatines is preserved; the fragment contacts the prefrontal via the prefrontal bar and appears to be heavily mediolaterally compressed, as the suture and the lateral edges form ridge-like structures that run anteroposteriorly along the palatines.

The posteriorly dislocated supraoccipital occupies less than a third of the width of the occipital surface and was not exposed on the skull roof in its original position. The posterior surface of the exoccipital is convex and subcylindrical dorsally, and flat and slightly concave ventrally, joining the basioccipital laterally and the basisphenoid posterolaterally. The basioccipital condyle is crescent shaped in posterior view and has a thick neck. The ventral margin of the basioccipital bears a W-shaped, ornamented margin which borders the median Eustachian foramen posteriorly, and extends dorsally to the posterior surface of the bone. The basisphenoid is exposed both on the ventral and the lateral side of the braincase and contacts the basioccipital and the exoccipital anteriorly and the quadrate medially. It bears a pair of posterolateral processes and a single small ventral process on both sides of the median Eustachian foramen. The laterosphenoid is subtriangular, borders the trigeminal foramen anteriorly, and bears a large anteroposterior ridge on each side, bordering a medial channel.

### Phylogenetic analyses

In order to rigorously assess the phylogenetic relationships of *Doratodon carcharidens*, we incorporated all available material of this taxon into updated versions of two separate global taxon-character datasets,each emphasizing a different taxonomic focus: one with enhanced neosuchian (Rummy et al. [30]) and the other with improved notosuchian sampling (Pinheiro et al. [31]) within Crocodyliformes (Supplementary Document). In addition, to test the taxonomic integrity of the materials referred to this taxon, we analyzed specimens both as separate and combined operational taxonomic units (OTU-s). In the ‘neosuchian’ dataset expanded from Rummy et al. [30] (Supplementary Document; Supplementary Data 1), the type locality (Campanian of Muthmannsdorf, Austria; ‘Doratodon_carcharidens_type’ OTU) and the Santonian Iharkút material (‘Doratodon_skull’ and ‘Doratodon_Ihar’ OTUs) are consistently recovered as sister taxa (Supplementary Document Figs. S5, S8) by the shared presence of a reduced antorbital fenestra (Ch. 67, state 1), ziphodont teeth (Ch. 120, state 1), a single laterally facing maxillary plane forming the external surface of the maxilla (Ch. 139, state 0), and a straight ventral maxillary edge (Ch. 183, state 0). The type locality material alone (Doratodon_carcharidens_type OTU) is consistently recovered as a neosuchian (Fig. S2).

In the dataset from Rummy et al. [30], the Iharkút and Muthmannsdorf type material of *Doratodon carcharidens*, either included separately or combined into a single OTU (Doratodon_carcharidens_type and Doratodon_skull OTUs together, Doratodon_S_T OTU, Doratodon_carcharidens_type and Doratodon_Ihar OTUs together, or Doratodon_ALL OTU), are consistently recovered as a member of the neosuchian clade Paralligatoridae based on a V-shaped intercondylar groove on the quadrate (Ch. 170 state 1) and a sharp ridge along the lateral side of the angular (Ch. 219, state 2). Within Paralligatoridae, *D. carcharidens* is placed in a basal position in a clade including *Paralligator* spp., *Yanjisuchus longshanensis, Turanosuchus aralensis*, *Kansajsuchus extensus*, the Dzharakuduk paralligatorid and *Scolomastax sahlsteini*. The inclusion of *D. carcharidens* into this clade is supported by the following synapomorphies: anteriorly tapering dentary symphysis (Ch. 154, state 0), the presence of an unsculpted region of the dentary below the tooth row (Ch. 155, state 1), weakly procumbent anterior dental alveoli (Ch. 262, state 1), the lack of a fossa at the anteromedial margin of the supratemporal fenestra (Ch. 265, state 1), and on some trees, a mediolaterally compressed and vertical dentary (Ch. 160, state 0). *Rugosuchus nonganensis, Shamosuchus djadochtaensis, Wannchampsus kirpachi*, and the Glen Rose Form (material likely representing *Wannchampsus* sp. as stated by Adams ^32^) are recovered as a separate paralligatorid clade that also includes *Tarsomordeo winkleri* on some trees, supported by two synapomorphies (see Supplementary Document). *Batrachomimus pastosbonensis* is not recovered in Paralligatoridae in any of our analyses. When the type material of *D. carcharidens* (lower jaw + maxilla) is excluded, the MTM PAL 2024.159.1 Iharkút skull alone (Doratodon_skull OTU) is recovered within Paralligatoridae in the same position as above, or in some trees as a basal mesoeucrocodylian sister to the fragmentary taxon *Sabresuchus sympiestodon*, the two sharing a midline ridge on the dorsal surface of the frontal and parietal (Char. 22, state 1) and an unsculpted lobe on the dorsolateral process of the squamosal (Char. 35, state 1; Figs. S3, S6).

In contrast to Pochat-Cottilloux et al. [33] and in spite of the addition of the taxon *Varanosuchus sakonnakhonensis*, our analyses do not recover Atoposauridae + Paralligatoridae as a clade unless the OTU used for *D. carcharidens* consists solely of the MTM PAL 2024.159.1 skull (Doratodon_skull), and even then only on some of the trees when the latter is placed in a basal mesoeucrocodylian position.

Using a Ziphosuchian-focused dataset (modified from Pinheiro et al. [29]; Supplementary Data 2), the type locality (Doratodon_carcharidens_type OTU) and Iharkút material (both Doratodon_skull and Doratodon_Ihar OTUs) of *D. carcharidens* consistently appear as sister taxa, sharing a single laterally facing maxillary plane forming the external surface of the maxilla (Ch. 128, state 0) (Fig. S12) and large foramina along the alveolar edge of the dentary (Ch. 342, state 1) (Fig. S15). United into a single OTU (Doratodon_S_T or Doratodon_ALL), *D. carcharidens* is recovered as the sister taxon of *Theriosuchus* sp. within Neosuchia (Fig. S14) supported by an exposed basisphenoid on the ventral surface of the braincase (Ch. 49, state 0), a midline ridge on the frontal and parietal (Ch. 20, state 1), a sharp ridge along the lateral surface of the angular (Ch. 205, state 2), and a posteriorly concave dentary with a single dorsal expansion (Ch. 148, state 2). Even when *D. carchariden*s is restricted to the type locality material (Doratodon_carcheridens_type OTU), it is still not consistently recovered as a ziphosuchian (Fig. S9).

*Ogresuchus furatus* is recovered as a neosuchian, either a member of Atoposauridae or a basal neosuchian sister to *Stolokrosuchus lapparenti,* with the expanded Rummy et al. [30] dataset and as an early diverging notosuchian with the Pinheiro et al. [31] dataset. There are only a few synapomorphies with either clade, usually the reduced contribution of the premaxilla to the narial border (Ch. 3, state 0 in Rummy et al. [30], Ch. 4, state 0 in Pinheiro et al. [31]) and the lateral edges of the suborbital bar being straight in the anterior portion (Ch. 278, state 0, Rummy et al. [30]), and flared posteriorly in the posterior portion (Ch. 279, state 1, Rummy et al. [30]; Ch. 260, state 1, Pinheiro et al. [31]).

## Discussion

### Doratodon carcharidens is a neosuchian

Previous phylogenies [5,20,28] and early systematic works [34] utilizing the then known incomplete material attributed to *Doratodon carcharidens* (mandible, maxilla, premaxilla and teeth) placed this taxon close to or within the Gondwanan ziphosuchian clade Sebecosuchia. The herein described partial skull referable to this taxon, however, unexpectedly but clearly implies a position within neosuchians regardless of taxon sampling. Within Neosuchia, a paralligatorid affinity (Figs. 6, S4-S8) is supported by two synapomorphies shared with the whole clade, and a further five synapomorphies shared with later diverging paralligatorids (see Results: Phylogenetic analyses). The referral of the Iharkút material to *D. carcharidens* is consistent with the phylogenetic analysis as it forms the sister taxon of the type material from Muthmannsdorf (Figs. S6, S9, S13, S16; see also Systematic Paleontology). A neosuchian placement is consistent with some early, non-phylogenetic systematic studies of this taxon [19,35].

**Fig. 5.:**
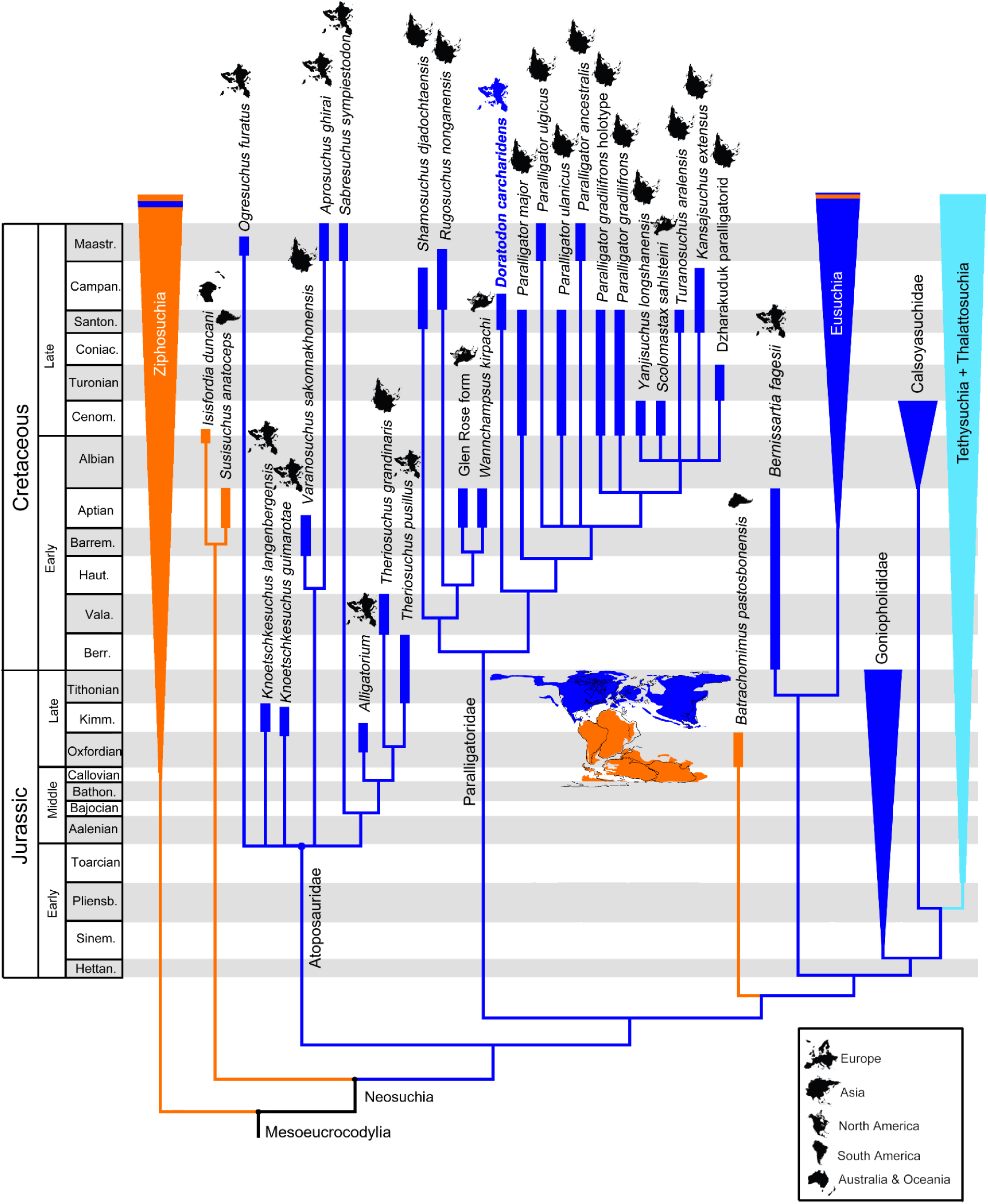
Time-calibrated reduced strict consensus tree resulting from the parsimony analysis of the expanded global dataset of Rummy et al. [30]. The *Doratodon carcharidens* terminal includes the type-locality and Iharkút materials. Other analyses incorporating terminals differentiated by locality or employing datasets with improved sampling of notosuchians [31] yield consistent results (Supplementary Document Figs. S4, S5, S7, S8, S11, S12, S14, S15). Gondwanan taxa are marked in orange, Laurasian taxa in dark blue, and aquatic clades with a global distribution in light blue. Paleogeographic map based on Scotese et al. [15], depicting the geography of the latest Berriasian. Temporal range data based on Turner [29], Rummy et al. [30], Pochat-Cottilloux et al. [33], Sellés et al. [38], Csiki-Sava et al. [42], and Dal Sasso et al. [89]. Abbreviations: Barrem., Barremian; Bathon., Bathonian; Berr., Berriasian; Campan., Campanian; Cenom., Cenomanian; Coniac., Coniacian; Haut., Hauterivian; Hettan., Hettangian; Kimm., Kimmeridgian; Maastr., Maastrichtian; Pliensb., Pliensbachian; Santon., Santonian; Sinem., Sinemurian; Vala., Valanginian.

**Fig. 6.**
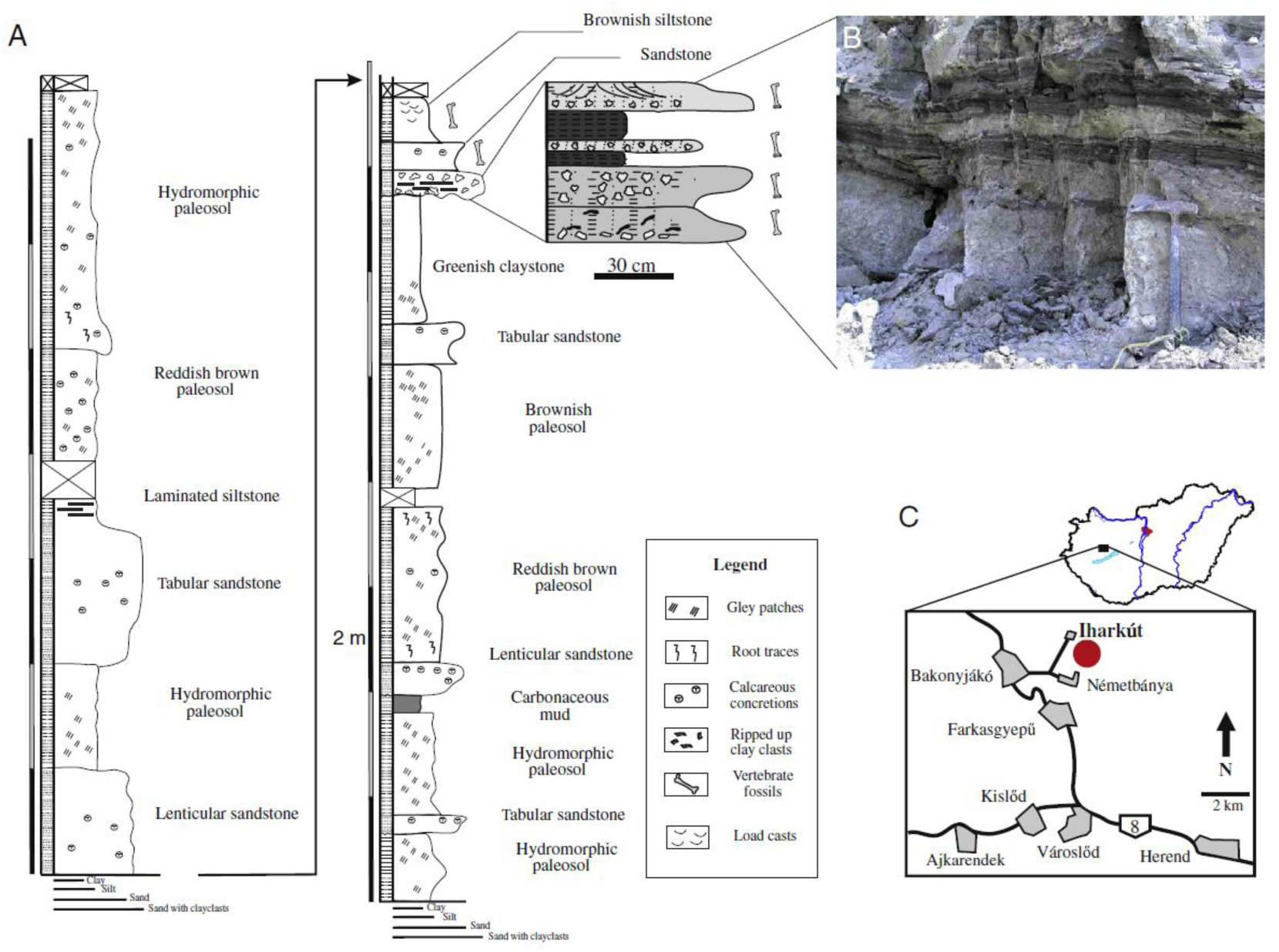
Geology and location of the Iharkút site. **A** Stratigraphic section of the Szál-6 outcrop where the specimen was found. **B** Photo of the lowermost bone-bearing unit exposed at the Szál-6 outcrop. **C** The location of the Iharkút locality in Western Hungary. Modified after Botfalvai et al. [85]

Some trees of the matrix with improved neosuchian sampling (Figs. S3, S6) place *D. carcharidens* in a basal position either within Neosuchia, or within Mesoeucrocodylia. In the latter case, it is recovered as the sister taxon of *Sabresuchus sympiestodon*, a fragmentary taxon of ambiguous phylogenetic position, recovered as an atoposaurid in our other analyses. However, all synapomorphies linking the two species together are widespread in both atoposaurids and paralligatorids, and characters recovered as plesiomorphic in *D. carcharidens* compared to mesoeucrocodylians or later branching neosuchians are also exhibited by derived paralligatorid taxa. As such, a basal neosuchian or mesoeucrocodylian position for either *Doratodon carcharidens* or *S. sympiestodon* might well be an artifact of missing data, as expanding the dataset with data from the type locality material of *D. carcharidens* consistently places the two taxa in Paralligatoridae and Atoposauridae respectively. *D. carcharidens* is known to co-occur with a species closely related to *Sabresuchus sympiestodon* [24], however, as discussed above, neither the MTM PAL 2024.159.1 skull nor the rest of the Iharkút *D. carcharidens* material can be referred to this taxon.

None of our analyses recover *D. carcharidens* consistently as a ziphosuchian, even though both matrices include several ziphosuchian taxa. In fact, a ziphosuchian placement is only supported in select trees when using an OTU consisting solely of the incomplete and fragmentary type locality material in the ziphosuchian-focused dataset of Pinheiro et al. [31]. Moreover, this placement is based on traits subject to ecomorphological convergence such as a ziphodont dentition, a mediolaterally compressed dentary and maxilla, and an anteriorly tapering dentary symphysis [36,37]. The complete *D. carcharidens* OTU is recovered as the sister taxon of *Theriosuchus* sp. in the Pinheiro et al. [31] dataset instead of the few paralligatorid taxa sampled by this matrix (Fig. S15). The difference in position from the Rummy et al. [30] dataset is therefore likely due to sampling and most of the synapomorphies are widespread in both paralligatorids and atoposaurids [33].

### *Ogresuchus furatus* is a possible atoposaurid

Our analyses of *Ogresuchus furatus* from the Late Cretaceous of Spain question the previously reported sebecid [38] or baurusuchian [39] affinities of this taxon and recover it either as an atoposaurid or basal neosuchian (Rummy et al. [30] dataset, Supplementary Document Figs. S2-S8) or an early diverging notosuchian (Pinheiro et al. [31] dataset, Supplementary Document Figs. S9-S15). One of the few synapomorphies uniting this taxon with notosuchians in the ziphosuchian-focused dataset of Pinheiro et al. [31] is the presence of prezygapophyses anteriorly exceeding the neural arch of the axis. However, this region is poorly known in most Mesozoic neosuchians from Europe and might have a broader distribution. The remaining synapomorphies placing this taxon with Ziphosuchia in these analyses, such as the anteriorly subparallel and posteriorly flared lateral edges of the suborbital bar and the reduced contribution of the premaxilla to the internarial bar, are homoplastic in the dataset within Mesoeucrocodylia and are furthermore present in *Theriosuchus* spp. and other atoposaurids [40], taxa that are known from the Cretaceous of Europe. The taxon sampling of Sellés et al. [38] is biased towards ziphosuchians even though the provenance, age and morphology of *O. furatus* is consistent with that of atoposaurid neosuchians [41].

A key trait linking *O. furatus* to Sebecidae in Sellés et al. [38] is a purported reduced maxillary dentition. Our direct observation of the specimen reveals, however, that the maxilla of the specimen (MCD-7149) is posteriorly incomplete: what Sellés et al. [38] interpret as the contact surface for the lacrimal is a damaged margin and cannot be conclusively identified as a suture (Fig. S2). Since the posterior process of the maxilla may have extended further and the number of alveoli may well have been higher, we code this character as unknown (Ch. 108 in the matrix derived from Pinheiro et al. [31]). Assuming that the maxilla of *O. furatus* is incomplete, it is consistent in proportions with *Sabresuchus sympiestodon* and related taxa from the Late Cretaceous of Europe in having a deep anterior half and a low posterior half, the latter containing the toothrow and extending below the jugal [24,41]. Although Sellés et al. [38] interpreted that the caniniform maxillary tooth is the 3rd in *O. furatus*, our direct observation of the maxilla revealed that it is anteriorly incomplete, and an additional alveolus may have been present, which would make the caninform tooth the 4th as in *Sabresuchus sympiestodon.* The incompleteness of the maxilla in *O. furatus* is furthermore supported by the extent of the maxillary palatal plate, which is still not terminating at the anteriormost alveolus as it would be otherwise expected. *O. furatus* shares several derived traits with *S. sympiestodon* and *Sabresuchus ibericus*, such as a pseudoziphodont teeth with apicobasal striae, and a slight depression on the ventral half of the lateral maxillary surface posterior to the caniniform tooth. Moreover, the dorsal and ventral half of the lateral maxillary surface of *O. furatus* are angled similarly as in *S. ibericus*, *S. sympiestodon* and *Knoetschkesuchus guimarotae* and the border between the two resembles the maxillary groove characteristic of *S. sympiestodon* [41]. In summary, we see no convincing evidence for the sebecid or ziphosuchian affinity of *O. furatus* and interpret this taxon as a neosuchian, most likely related to *Sabresuchus spp.*, taxa that are widely distributed in Europe.

#### Implications for Mesozoic European paleogeography

Rabi & Sebők [5], in light of the then available more incomplete material, discussed the implication of the sebecosuchian affinities of *Doratodon carcharidens* for the establishment of faunal contact between Africa and Europe during the Late Cretaceous as early as the Santonian. However, the more comprehensive phylogenetic analyses of the present study, including the herein described partial skull, of *Doratodon carcharidens*, consistently find this taxon closely related to representatives of Paralligatoridae (Fig. 5). a clade hitherto almost exclusively known from the North American and Asian Cretaceous [29]. The only putative paralligatorid outside Laurasia is *Batrachomimus pastosbonensis* from the Upper Jurassic of Brazil [30, 43], a taxon placed outside this clade in both of our phylogenetic analyses (Fig. 5; Supplementary Document S1.2). The new, more complete material of *Doratodon carcharidens* integrated into global phylogenies therefore reinterprets this taxon as of Laurasian origin. The earliest unambiguous paralligatorids come from the Lower Cretaceous (Aptian) of North America but the geographic origin of the clade within Laurasia remains uncertain as the ingroup phylogeny is far from being settled and sampling is likely biased [29,30,32,44,45]. Literal reading of the fossil record would imply North American origin and subsequent dispersal to Asia via Beringia [46,47]. The spatiotemporally closest paralligatorids to *D. carcharidens* suggest dispersal to Europe from central Asia. These hypotheses, however, lack consistent support from phylogenies. Moreover, a vast epicontinental sea, the Turgai Strait, separated Europe from Asia during much of the Mesozoic and Cenozoic-likely acting as a significant barrier for faunal dispersal [42,48,49]. Alternatively, *D. carcharidens* represents an Euramerican relict that is still yet to be sampled in the poorly known Early Cretaceous of Europe. Remarkably, Ősi et al. [50] reported a crocodyliform tooth from the Albian of western Hungary similar in morphology to those of the North American Aptian taxon, *Wannchampsus kirpachi*, and suggested the possible early presence of paralligatorids in Europe.

Another crocodyliform in the Late Cretaceous of Europe with a purported Gondwanan link is *Ogresuchus furatus* from the Maastrichtian of Spain. This taxon has been considered the oldest representative of Sebecidae and used as evidence for a European origin of sebecids with subsequent dispersal to South America (via Africa [38]). The present study, however, demonstrates that this taxon is most likely a representative of a Laurasian clade and closely related to the atoposaurid-like neosuchian *Sabresuchus sympiestodon* (Fig. 5). With the exception of *Isisfordia duncani* and *Susisuchus anatoceps*, our phylogenetic results tentatively support a scenario in which the divergence between ziphosuchian and neosuchian crocodyliforms may have been influenced by the initial breakup of Pangaea into Gondwana and Laurasia (Fig. 5).

During the Late Cretaceous, Europe was a tectonically active archipelago of ephemeral islands, with the ‘Iharkút landmass’, as part of the Austro-Alpian block, existing for about 4-6 million years from the late Turonian until the middle Santonian [42]. The Late Cretaceous fauna assemblages of Europe are considered to consist of endemic relic clades as well as immigrants, particularly from Gondwana [3,5,6,10,42]. Based on the inferred Gondwanan origin of numerous tetrapod clades, previous studies have proposed a relatively late separation of Europe from Gondwana and associated prolonged retention of faunal connections throughout the Cretaceous period (Eurogondwana and Atlantogea models). These hypotheses inherently rely on our understanding of phylogenetic relationships of extinct taxa with predominantly poor preservation potentials, i.e., terrestrial and freshwater vertebrates. As the case of *Doratodon carcharidens* aptly highlights, phylogenetic inference is strongly subject to fossil completeness as well as taxon sampling. Moreover, the Mesozoic continental record of Gondwana and Europe is highly incomplete compared to North America and Asia. We argue that, in order to infer faunal connections between Europe and Gondwana, an extinct taxon must satisfy a pair of key criteria: (i) a well-supported sister-relationship between taxa from both landmasses, and (ii) a distribution that is not cosmopolitan. Taxa employed to hypothesize dispersal events or an early Eurogondwanan distribution should be robust to alternative phylogenetic placements and to the possibility that they represent vicariant descendants of a previously widespread clade. Scrutinizing previously used evidence reveals that only a single taxon fulfills both of these criteria. Besides crocodyliforms, several other taxa from the Late Cretaceous of Europe (including the Iharkút), have been proposed to represent Gondwanan dispersals (Rabi & Sebők [5] and references therein). Among these, bothremydid turtles are known by far from the most complete material, including near-complete skeletons. Rabi et al. [51] proposed an African origin of European bothremydids but subsequent fossil finds and phylogenies raised the strong possibility of a dispersal from North America [52, 53]. While no doubt an originally Gondwanan clade [52,54], their wide Late Cretaceous geographic distribution and partially unresolved ingroup relationships currently hinders reconstructing the biogeographic history of European bothremydids, especially considering that most if not all bothremydids were saltwater tolerant [51,54,55]. Other pan-pleurodiran turtles present in the Cretaceous of Europe include representatives of Dortokidae [56,57], an incompletely known group best interpreted as stem-pleurodires and so far considered endemic to Europe [58]. While Pan-Pleurodira is Gondwanan in origin [52], some stem-pleurodires are considered saltwater tolerant, which likely facilitated their dispersal to Europe at latest by the Late Jurassic [59]. Dortokids cannot be traced back beyond the Early Cretaceous [60,61] and phylogenies, although based on limited character evidence, place them in a more crownward position [58]. Even if dortokids were not descendants of Late Jurassic European taxa, they may have dispersed to Europe via marine routes since both platychelyid stem-pleurodires and several crown-pleurodires appear to be saltwater tolerant [54,57,59,62]. Shell histology has been used to infer freshwater habitat for Dortokidae [63] but this is not incompatible with saltwater tolerance.

Phylogenies of Cretaceous macronarian sauropod dinosaurs from Europe often recover these taxa near or within Gondwanan clades [14,23,64,65,66]. The case of *Doratodon carcharidens* presented here, however, demonstrates that a literal interpretation of these cladograms as evidence for faunal connections can be highly misleading. The relevant sauropod record supporting the phylogenies in question is spatiotemporally highly uneven in sampling as well as extremely fragmentary, consisting of postcranial elements, most often vertebrae. In addition, the proposed relationships have a poor stratigraphic fit and imply overly complex biogeographic histories, including multiple dispersals involving Asia and Australia besides Europe, Africa and South America. Similar arguments can be made against the purported evidence provided by European abelisaurid theropods [4,67,68,69] with often uncertain phylogenetic relationships [70,71] based on poorly diagnostic material [72,73] with a long unsampled history between the Late Jurassic and the Late Cretaceous [74]. Spinosaurid theropods have been hypothesized to originate in Europe and disperse to Africa in the Late Cretaceous but the supporting phylogeny relies on highly fragmentary taxa (*Vallibonavenatrix* and *Camarillasaurus* as early diverging spinosaurids [10]). In addition to phylogenetic uncertainty, we further argue that the markedly incomplete sampling of Mesozoic European dinosaurs, in conjunction with the widespread geographic distribution of their more inclusive clades, prevents the definitive exclusion of the hypothesis that Cretaceous taxa represent relict lineages persisting from pre-Pangaean fragmentation times. Consequently, vicariant speciation remains a plausible alternative to intercontinental dispersal in explaining the biogeographic distribution of these taxa. Vicariant speciation is therefore a viable alternative to intercontinental dispersal when explaining the distribution of these taxa. We therefore find it premature to use the current Cretaceous dinosaur record for inferring faunal and land connections between Europe and Gondwana. Our caution is in line with a recent review highlighting the global distribution of multiple dinosaur lineages, including macronarian sauropods, before giving rise to Cretaceous faunas [75]. A notable exception may be arenysaurinine hadrosaurids, a group so far only reported from the Maastrichtian of Iberia and Morocco [11,12]. The restricted distribution and origin of this group remains to be explained but their cranial anatomy is relatively better known compared to other ‘Gondwanan’ taxa from Europe. Hadrosaurids being an entirely Late Cretaceous clade, it is unlikely that arenysaurinines had considerably earlier unsampled presence in Europe and hadrosaurids are otherwise completely absent from the African record. The presence of arenysaurinines in Morocco is substantiated with a number of cranial characters and has been explained with oceanic dispersal from Iberia [11,12].

Other European taxa previously discussed in the context of possible ‘Gondwanan’ clades include neobatrachian anurans [76,77,78] and pan-lepisosteid fishes [79,80,81], both of which are known from highly fragmentary remains. However, these interpretations have been subsequently challenged due to limited and potentially convergent morphological character evidence [77], as well as conflicting phylogenetic placements in the case of neobatrachians [78]. For pan-lepisosteids, the absence of phylogenetic support [82,83], combined with their widespread Cretaceous distribution and inferred tolerance for saltwater [83], renders a dispersal scenario from North America a plausible alternative explanation.

## Conclusions

The European Late Cretaceous crocodyliform *Doratodon carcharidens* is reinterpreted as a representative of the Laurasian neosuchian clade Paralligatoridae that evolved convergent ecomorphological traits with Gondwanan ziphosuchian sebecosuchians including ziphodont dentition and an oreinirostral snout. Our analyses recover the only putative Gondwanan paralligatorid, *Batrachomimus pastosbonensis* outside of this clade, further reinforcing the Laurasian distribution and origin of *D. carcharidens* and other paralligatorids. In addition, we reinterpret *Ogresuchus furatus*, another Late Cretaceous European taxon previously considered a ziphosuchian, as related to the Eurasian neosuchian clade Atoposauridae. The case of *Doratodon carcharidens*, a taxon whose previously proposed Gondwanan origin has been refuted through the analysis of more complete fossil material, underscores the importance of re-evaluating biogeographic and systematic interpretations based on fragmentary remains in other clades. Our critical reassessment reveals that previous faunal evidence for connections between Europe and Africa during the Cretaceous is ambiguous by relying on stratigraphically restricted and incompletely preserved fossil records and conflicting or poorly supported phylogenetic relationships. It remains plausible that taxa previously interpreted as indicators of faunal and terrestrial connectivity instead represent vicariant Pangean relicts. Notable exceptions likely include arenysaurinine hadrosaurids that are best explained by a latest Cretaceous dispersal from Europe to Africa. In conclusion, we advocate a reconsideration of the traditional Laurasia– Gondwana dichotomy, given the lack of compelling evidence supporting the Eurogondwana model.

## Methods

The partial skull MTM PAL 2024.159.1 described here comes from the Iharkút vertebrate site and was recovered in 2018 except for the associated right maxilla (belonging to the same individual) recovered in 2024. An isolated ectopterygoid likely but not unambiguously belonging to this skull was furthermore found in 2024. Other cranial material from the site referred to *Doratodon carcharidens* includes an isolated quadrate (MTM PAL 2013.67.1.), pterygoid (MTM PAL 2013.64.1.); and material described in Rabi & Sebők [5]: right premaxilla (MTM PAL 2014.122.1), fragmentary left maxilla (MTM PAL 2013.65.1), fragmentary left dentary (MTM V2010.237.1) and a fragmentary right dentary (MTM PAL 2013.66.1).

The type-locality material of *Doratodon carcharidens* is from the lower Campanian Grünbach Formation of Austria and consists of a mandible (holotype, IPUW2349/57), a partial maxilla (IPUW2349/5) and possibly a parietal (UWPI 2349/54) from the University of Vienna’s paleontological collection. We furthermore studied the holotype of *Ogresuchus furatus* (MCD-7149) [38].

### Phylogenetic analysis

We included *Doratodon carcharidens* in the taxon-character matrix of Rummy et al. [30] densely sampling neosuchians (Supplementary Data 1), adding four taxa from Pochat-Cottilloux et al. [33] in addition to five OTUs of *Doratodon carcharidens*, and the crocodyliform *Ogresuchus furatus* from the Late Cretaceous of Spain, previously considered closely related to Gondwanan crocodyliforms. The OTUs are described below. We furthermore tested the placement of *D. carcharidens* in a separate dataset with a ziphosuchian focus (Pinheiro et al. [31]) and added *O. furatus* and the five *D. carcharidens* OTUs to the previous matrix (Supplementary Data 2). Character 138 of Rummy et al. [30] and Character 127 of Pinheiro et al. [31] were reset as inactive in our analyses, in addition to the inactive characters in Rummy et al. [30] and prior works (see Supplementary Materials and Supplementary Methods). We updated several character scores for *Batrachomimus pastosbonensis, Sabresuchus sympiestodon* and other taxa (see Supplementary Document: Dataset based on Rummy et al. [30], Dataset based on Pinheiro et al. [31]).

We tested the phylogenetic position of *Doratodon carcharidens* using different operational taxonomic units (OTUs): *i)* partial skull (MTM PAL 2024.159.1) from the Santonian of Iharkút (Doratodon_skull); *ii)* type locality material from the lower Campanian Grünbach Formation of Austria, including the mandible IPUW 2349/57 and maxilla IPUW2349/5 (Doratodon_carcharidens_type); *iii)* the MTM PAL 2024.159.1 skull and the type locality material coded as a single OTU (Doratodon_S_T); *iv)* all the material from Iharkút including the MTM PAL 2024.159.1 skull, isolated quadrate (MTM PAL 2013.67.1.), isolated pterygoid (MTM PAL 2013.64.1.), and the material described in Rabi & Sebők [5] (Doratodon_Ihar); *v)* and the Iharkút and type locality material as a single OTU (Doratodon_ALL) (see Supplementary Methods).

A heuristic search was performed on the dataset modified from Rummy et al. [30] in TNTv.1.6 [84], performing 1000 replications of the tree-bisection-reconnection algorithm with 10 trees in each; or using the xmult=hits30 command (Goloboff et al. 2008) to achieve at least 30 replications with maximum-parsimony trees (MPTs) in cases where the analysis failed to produce at least 60 MPTs. For the matrix based on Pinheiro et al. [31], we followed the method described therein (see Supplementary Document: Dataset based on Pinheiro et al. [31]). Detailed descriptions and consensus trees resulting from our analyses, as well as any changes made to the matrices and sources for the included taxa are available in the Supplementary Document.

### Institutional abbreviations

IPUW/UWPI, Institute of Paleontology, University of Vienna, Austria; MCD, Museu de la Conca Dellà, Isona, Spain; MTM, Hungarian Natural History Museum, Budapest, Hungary.

### Geological setting

The Iharkút vertebrate site (Fig. 6.) is located near the villages of Bakonyjákó and Németbánya, within a former open pit bauxite mine. The fossiliferous strata exposed at the Iharkút site belong to the Santonian [25] Csehbánya Formation, overlaying the Upper Cretaceous Nagytárkány Bauxite, which was deposited in the karstic depressions of the Upper Triassic Main Dolomite Formation [85]. The Csehbánya Formation consists of terrestrial siliciclastic sediments of cyclically changing grain size, periodically interrupted by coal seams. Botfalvai et al. [86] identified four facies associations from the Iharkút locality, the first consisting of heterolithic fluvial channels and flash flood deposits (the latter of which being the most important fossiliferous strata), and the other three being floodplain deposits with plant remains, mollusc- and microfossil-rich clayey shallow lake deposits, and paleosols. The partial skull here described (MTM PAL 2024.159.1) was found at site Szál-6, which exposes sequences of clastic breccias with erosional boundaries overlaid by laminar organic-rich siltstone with plant remains, and sandstones, all three of which are fossiliferous. The skull itself was recovered positioned on the boundary between the underlying greenish claystone and the clastic breccia layer. Faunal elements reported from the locality include lepisosteiform and pycnodontiform fish, albanerpetontid and anuran amphibians, turtles, squamates, ornithischian, sauropod and theropod dinosaurs (including birds), pterosaurs and crocodyliforms [24,87]. Crocodyliforms include the heterodont hylaeochampsid *Iharkutosuchus makadii* [88], the allodaposuchid *Allodaposuchus* sp. [24], a taxon closely related to *Theriosuchus (Sabresuchus) sympiestodon* [24], and the ziphodont *Doratodon carcharidens* [5].

## Author contributions

A.Ő. directed the collection of the specimens and provided geological and faunistic context. M.Sz. described the specimens, performed the phylogenetic analyses and reviewed the European record of relevant clades. M.R. conducted the paleobiogeographic interpretations. All authors designed the study, collected data, discussed the results, and wrote and reviewed the manuscript.

## Data availability

All data generated or analysed during this study are included in this published article and its supplementary information files.

## Acknowledgements

We thank Martin Maslo, Sebastian Stumpf (University of Vienna), Márton Szabó, Piroska Pazonyi (Hungarian Natural History Museum), Àngel Galobart, Irina Fernández, Laura Fàbrega, and Óscar Castillo-Visa (Institut Català de Paleontologia Miquel Crusafont) for their help in accessing and examining the specimens in their care. We are thankful to Albert G. Sellés (Institut Català de Paleontologia Miquel Crusafont), Aline Marcele Ghilardi (Universidade Federal do Rio Grando do Norde), André Eduardo Piacentini Pinheiro (Universidade do Estado do Rio de Janeiro), Paul Rummy (Chinese Academy of Sciences), Rafael Delcourt (Universidade Estadual de Campinas), and Yohan Pochat-Cottilloux (Université Claude Bernard Lyon 1) for sharing their research, and to the Willi Hennig Society for granting free access to TnT. We are thankful to Nóra Sebők for her illustrations of the isolated specimens described above, and János Magyar for his technical assistance and practical advice. Our research would not have been possible without the members of the Iharkút expeditions from 2000 to 2024; special thanks are in order to Balázs Sütöri and Emma Ayres for finding and recovering the new specimens examined in this study.

The fieldwork and research of the Iharkút site was supported by the Hungarian Academy of Sciences, the Hungarian Natural History Museum, the National Geographic Society, the Hungarian Scientific Research Fund, the Eötvös Loránd University Departments of Paleontology and Applied and Physical Geology, the Jurassic Foundation, the Hantken Miksa Foundation, the Hungarian Dinosaur Foundation, the Pro Renovanda Cultura Hungariae Foundation, Nemzeti Kutatási és Technológiai Hivatal, and the Hungarian Oil and Gas Company (MOL).

