## Supplementary Document for "Cretaceous crocodyliform reconciles conflicting evidence on the Mesozoic paleogeography of Europe during the Gondwana-Laurasia split"

**Table of contents**

Supplementary Materials	2

Sources for taxa	2

Institutional abbreviations	3

Supplementary Methods	4

Remarks	4

Doratodon OTUs used throughout phylogenetic analyses	5

Dataset based on Rummy et al.	7

Analyses in the Rummy et al. dataset	9

Dataset based on Pinheiro et al.	28

Analyses in the Pinheiro et al. dataset	29

Supplementary References	43

### Supplementary Materials

#### Sources for taxa

*Acynodon adriaticus*: Delfino et al.^1^, MCSNT 57248 (Text-Fig. 3)

*Alligator mississippiensis:* Pochat-Cottilloux et al.^2^ phylogenetic matrix

*Allodaposuchus precedens:* Martin^3^, PSMUBB V438 (Fig. 4B)

*Aprosuchus ghirai*: Venczel & Codrea^4^, UBB V.562/1 (Fig. 6); Pochat-Cottilloux et al.^2^ phylogenetic matrix

*Asiatosuchus germanicus:* Vasse^5^, ISS 1 a. from the Laboratoire de Paléontologie des Vertébrés et de Paléontologie Humaine, Paris (Plate 1)

*Batrachomimus pastosbonensis*: Montefeltro et al.^6^, LPRP/USP-0617 (Figs. 2-3)

*Bernissartia fagesii:* Pochat-Cottilloux et al.^2^ phylogenetic matrix

*Calsoyasuchus valliceps*: Tykoski et al.^7^, TMM 43631-1 (Fig. 1)

*Crocodylus niloticus*: Shaker & El-Bably^8^, specimen number not given (Fig. 3)

*Dakosaurus andiniensis*: Pol & Gasparini^9^, MOZ 6146P (Fig. 2)

*Diplocynodon hantoniensis:* Rio et al.^10^, NHMUK OR 30393 (Fig. 4)

*Doratodon carcharidens*, Iharkút material: direct observation of MTM PAL 2024.159.1 (partial skull), MTM PAL 2013.67.1. (quadrate), MTM PAL 2013.64.1. (pterygoid); Rabi & Sebők^11^ and direct observation of MTM PAL 2014.122.1(fragmentary premaxilla), MTM PAL 2013.65.1(fragmentary left maxilla), MTM V2010.237.1(fragmentary left dentary), MTM PAL 2013.66.1(fragmentary right dentary).

*Doratodon carcharidens*, type locality (Muthmannsdorf) material: direct observation of IPUW2349/57 (mandible, holotype) and IPUW2349/5 (partial maxilla)

*Gavialis gangeticus*: Gold^12^, YPM HERR-008438, TMM-M-5490 (Figs. 8, 19)

*Goniopholis baryglyphaeus:* De Andrade et al.^13^, MG ‘Gui Croc 1’ (Fig. 5)

*Goniopholis simus:* De Andrade et al.^13^, BMNH 41098 (Fig. 3)

*Kansajsuchus extensus:* Kuzmin et al.^14^, PIN 2399/301, PIN 2399/445 (Figs. 1-3)

*Kaprosuchus saharicus:* Sereno & Larsson^15^, MNN IGU12 (Figs. 33, 34, 36B)

*Knoetschkesuchus guimarotae:* Schwarz & Salisbury^16^, IPFUB Gui Croc 7308-1, IPFUB Gui Croc 7309 (Figs. 2-3); Pochat-Cottilloux et al.^2^ phylogenetic matrix

*Knoetschkesuchus langenbergensis:* Schwarz et al.^17^, DFMMh/FV 200 (Fig. 4), DFMMh/FV 605 (Fig. 5); Pochat-Cottilloux et al. (2024) phylogenetic matrix

*Lomasuchus palpebrosus:* Gasparini et al.^18^ (1991), MOZ 4084 PV, (Figs. 3, 5)

*Ogresuchus furatus*: Sellés et al.^19^, direct observation of MCD-7149 (holotype)

*Paluxysuchus newmani:* Adams^20^ (2013), SMU 76601 (Fig. 3C)

*Paralligator tersus:* Turner^21^, PIN 3141-501 (Fig. 11C)

*Paralligator ulgicus:* Turner^21^, PIN 3458/501 (Fig. 9)

*Sabresuchus sympiestodon*: Martin et al.^22^, LPB (FGGUB) R.1781, MCDRD 793, LPB (FGGUB)

R.1945 (Figs. 1, 2, 4, 5)

*Shamosuchus djadochtaensis:* Turner^21^, AMNH FARB 6412 (Fig. 2), IGM 100/1195 (Fig.3)

*Theriosuchus grandinaris*: Pochat-Cottilloux et al.^2^ phylogenetic matrix

*Theriosuchus pusillus*: photographs of NHMUK 48227-8, 48282, 48282.a, and 48240.a by Rabi M.

*Varanosuchus sakonnakhonensis:* Pochat-Cottilloux et al.^2^ phylogenetic matrix; “V_sakonnakhonensis_h” is based on their coding for the holotype SM-2021-1-97/101 and “V_sakonnakhonensis_r” is based on their coding for the referred specimen SM-2023-1-16.

#### Institutional abbreviations

**AMNH FARB,** American Museum of Natural History, Collection of Fossil Reptiles, Amphibians, and Birds, New York, USA; **IGM,** Mongolian Institute of Geology, Ulaanbaatar, Mongolia; **IPUW**: Institute of Paleontology, University of Vienna, Austria; **LPB (FGGUB)**, Laboratory of Paleontology, Faculty of Geology and Geophysics, University of Bucharest, Bucharest, Romania; **LPRP/USP**, Laboratório de Paleontologia de Ribeirão Preto-USP, Ribeirão Preto, Brazil; **MCD**, Museu de la Conca Dellà, Isona, Spain; **MCDRD**, Muzeul Civilizației Dacice și Romane, Deva, Romania; **MCSNT**, Museo Civico di Storia Naturale di Trieste; **MG**, Museu Geológico, Lisboa, Portugal; **MNN**, Muséum National du Niger, Niamey, République de Niger; **MOZ**, Museo Profesor J. Olsacher, Zapala, Neuquén Province, Argentina; **MTM**, Hungarian Natural History Museum, Budapest, Hungary; **NHMUK**, Natural History Museum, London, UK; **PIN**, Borissiak Paleontological Institute, Russian Academy of Sciences, Moscow, Russia; **PSMUBB**, Paleontology-Stratigraphy Museum, University Babes-Bolyai, Cluj-Napoca, Romania; **SM**, Sirindhorn Museum, Kalasin, Thailand; **SMU**, Southern Methodist University Shuler Museum of Paleontology, Dallas, Texas, U.S.A.; **TMM**, Texas Memorial Museum, Austin, Texas, U.S.A.; **UBB**, Babeș-Bolyai University, Cluj-Napoca, Romania.

### Supplementary Methods

#### Remarks

In our phylogenetic analyses, we used a neosuchian-focused dataset from Rummy et al.^23^, and a ziphosuchian-focused dataset from Pinheiro et al.^24^, to account for the bias in character sampling^25,26^ present in each matrix. We used Mesquite version 3.81^27^ to edit the taxon-character matrices, and performed phylogenetic analyses on them using the software Tree analysis using New Technology (TnT 1.6^28^). We employed several OTUs based on various specimens attributed to *D. carcharidens* (see below).

#### *Doratodon* OTUs used throughout the phylogenetic analyses

**Doratodon_skull**: MTM PAL 2024.159.1 (partial skull from the Iharkút locality)

**Doratodon_carcharidens_type**: IPUW2349/57 (mandible, holotype), IPUW2349/5 (partial maxilla), both from the Muthmannsdorf type locality.

**Doratodon_Ihar**: all of the material attributed to *D. carcharidens* from the Iharkút locality: MTM PAL 2024.159.1 (partial skull), MTM PAL 2013.67.1. (quadrate), MTM PAL 2013.64.1. (pterygoid) (Fig. S1); MTM PAL 2014.122.1(fragmentary premaxilla), MTM PAL 2013.65.1(fragmentary left maxilla), MTM V2010.237.1(fragmentary left dentary), MTM PAL 2013.66.1(fragmentary right dentary).

**Doratodon_S_T**: MTM PAL 2024.159.1 (partial skull from the Iharkút locality), IPUW2349/57 (mandible, holotype from the Muthmannsdorf locality), IPUW2349/5 (partial maxilla from the Muthmannsdorf locality)

**Doratodon_ALL**: all of the material included in Doratodon_Ihar and Doratodon_carcharidens_type.

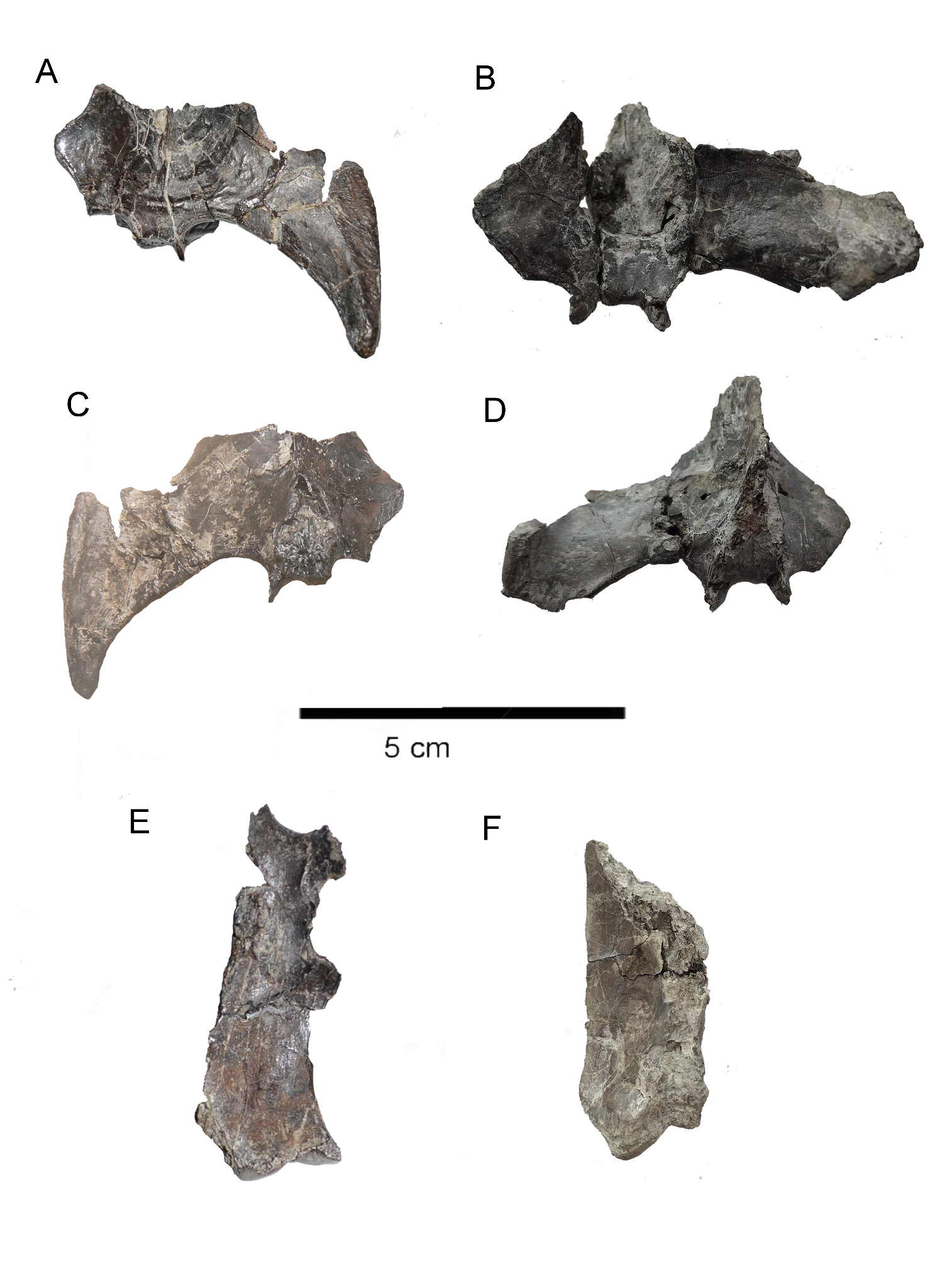

Fig. S1: Comparison of isolated cranial elements assigned to *D. carcharidens* from the Santonian Iharkút locality with the equivalent elements of the MTM PAL 2024.159.1 skull. A. ventral view of MTM PAL 2013.64.1.; B. ventral view of the pterygoid belonging to the MTM PAL 2024.159.1 skull. C. dorsal view of MTM PAL 2013.64.1.; D. dorsal view of the pterygoid belonging to the MTM PAL 2024.159.1 skull. E. ventral view of MTM PAL 2013.67.1. F. ventral view of the isolated posterior quadrate belonging to the MTM PAL 2024.159.1 skull.

### Dataset based on Rummy et al.

We used the neosuchian-focused dataset from Rummy et al.^23^, based on Kuzmin et al.^14^ and Turner^21^, with the addition of the aforementioned *Doratodon* OTUs, *Ogresuchus furatus*, *Aprosuchus ghirai*, *Theriosuchus grandinaris*, *Knoetschkesuchus langenbergensis* and two OTUs of *Varanosuchus sakonnakhonensis*. In addition, we updated the coding for *Knoetschkesuchus guimarotae* based on that of Pochat-Cottilloux et al.^2^, and edited the scoring for Characters 22 (0→1) and 67 (3→?) for *Sabresuchus sympiestodon,* and Characters 168 (1→?) and 288 (1→0) for *Batrachomimus pastosbonensis* (see Sources for taxa).

As in Rummy et al.^23^ and prior works, we used *Gracilisuchus stipanicicorum* as the outgroup taxon, and set Characters 5, 277 and 281 as inactive. We also set Character 138 (“Large and aligned neurovascular foramina along the lateral maxillary surface: absent (0), present (1)”) as inactive, due to inconsistencies in its coding in this matrix and previous works. Originally, these foramina were coded as present only in notosuchian taxa and *Iharkutosuchus makadii*, even though the foramina are clearly visible (in some cases, explicitly marked in figures or mentioned in the description of specimens) in several neosuchian and non-notosuchian ziphosuchian taxa including *Theriosuchus pusillus*, *Knoetschkesuchus* spp., *Kansajsuchus extensus*, *Lomasuchus palpebrosus*, *Kaprosuchus saharicus*, *Dakosaurus andiniensis*, *Paluxysuchus newmani*, *Goniopholis* spp., *Calsoyasuchus valliceps*, *Acynodon adriaticus*, *Allodaposuchus precedens*, *Gavialis gangeticus, Crocodylus niloticus,* *Asiatosuchus germanicus*, *Diplocynodon hantoniensis*, *Alligator mississippiensis, Bernissartia fagesii*, *Shamosuchus djadochtaensis, Paralligator ulgicus, Paralligator tersus,* and *Sabresuchus sympiestodon* (see Sources for taxa). In addition, the difference in coding cannot be attributed to a size threshold (despite the definition of the character stating ‘large’), as the size of these foramina differs among taxa coded as ‘present’, e.g. in the case of *Araripesuchus tsangatsangana* versus *Simosuchus clarki* ^29,30^. While we have recoded this character for the taxa listed above, a complete, consistent recoding for all taxa has proven impossible, as we did not have access to specimens or good quality photographs of several key taxa (e.g. the ‘Glen Rose Form’, *Paralligator major, Paralligator gradilifrons, Paralligator ulanicus*) to clearly determine the distribution of these foramina among Mesoeucrocodylia. As such, either leaving the character as is, or using the recoded version without accounting for all taxa would introduce bias into our analyses. In addition, these foramina are widespread among tetrapods (e.g. ^31,32^), making their relevance to mesoeucrocodylian evolution uncertain.

For analyses using the neosuchian-focused dataset, we used TNT’s Traditional Search option, with memory space for 10000 trees to perform 1000 replications of the tree-bisection-reconnection algorithm with 10 trees saved from each, and keeping the most parsimonious trees (MPTs) in the end; or alternatively using the xmult=hits30 command to run complex analyses (see <https://isu-molphyl.github.io/EEOB563/computer_labs/lab2/TNT.html>) until maximum parsimony has been reached 30 times if the previous method failed to produce at least 60 MPTs.

#### Analyses in the Rummy et al. dataset

**Analysis with Doratodon_carcharidens_type OTU (Fig. S2)**

The analysis of the type locality material recovered 65 trees, with a length of 1774. *Doratodon carcharidens* is recovered as the sister taxon of *Calsoyasuchus valliceps* in a derived position within Goniopholididae, on the basis of a shared antorbital fenestra.

*Ogresuchus furatus* is recovered as either a basal neosuchian and the sister taxon of *Stolokrosuchus lapparenti* with which it shares a notch in the premaxilla on the lateral edge of the external nares and a posteriorly flared suborbital bar; or an atoposaurid in a clade with *Sabresuchus sympiestodon*, *Alligatorium* and *Theriosuchus* spp., with which it shares an anteriorly straight suborbital bar. *Batrachomimus pastosbonensis* is recovered as an indeterminate neosuchian more derived than Atoposauridae, but more basal than all other neosuchians. *Sabresuchus sympiestodon* is recovered as the sister taxon of the clade comprising *Alligatorium* and *Theriosuchus* spp. within Atoposauridae.

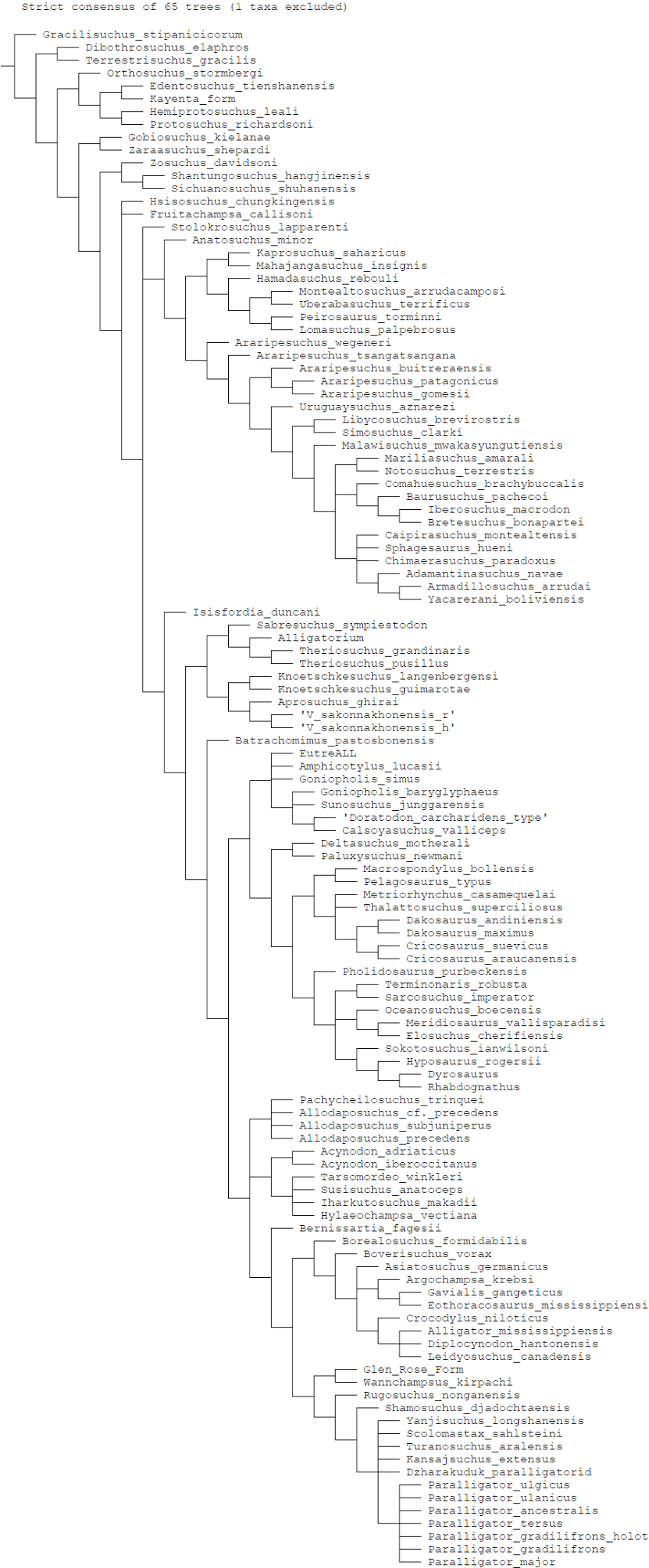

Fig. S2: Strict consensus of 65 trees from the analysis of the *Doratodon carcharidens* type locality (Muthmannsdorf) material in the first matrix, with *Ogresuchus furatus* pruned.

**Analysis with MTM PAL 2024.159.1 skull OTU (Fig. S3)**

Our first analysis of the MTM PAL 2024.159.1 skull recovered 80 trees, with a length of 1775. *Doratodon* is recovered as either a basal mesoeucrocodylian and the sister taxon of *Sabresuchus sympiestodon*, with which it shares a midline ridge on the dorsal surface of the frontal and parietal (Ch. 22, state 1) and an unsculpted lobe on the dorsolateral process of the squamosal (Ch. 35, state 1), or a basal neosuchian based on a horizontally projected posterior squamosal process (Ch. 36, state 0), and the lateral exposure of the basisphenoid on the braincase (Ch. 147, state 1), with *S. sympiestodon* being the sister taxon of the clade formed by *Alligatorium* and *Theriosuchus* spp. within Atoposauridae. *Ogresuchus furatus* is recovered as either an indeterminate atoposaurid based on the subparallel margins of the anterior half of the suborbital fenestra (Ch. 278, state 0), or a basal neosuchian and the sister taxon of *Stolokrosuchus lapparenti* based on a notch in the premaxilla on the lateral edge of the external nares (Ch. 123, state 1) and a posteriorly flared suborbital bar (Ch. 279, state 1).

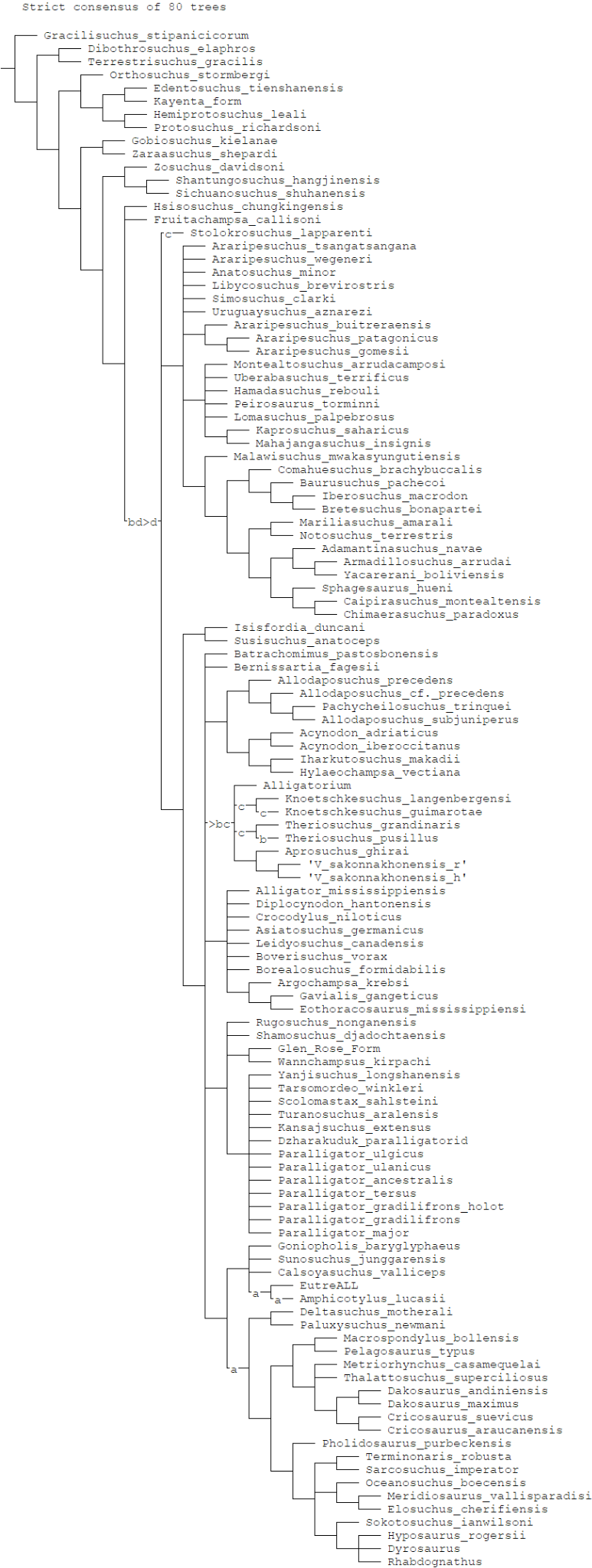

Fig. S3: Strict consensus of 80 trees from the analysis of the MTM PAL 2024.159.1 skull in the first matrix, with the possible positions of the following taxa: a. *Goniopholis simus*, b. *Sabresuchus sympiestodon*, c. *Ogresuchus furatus*, d. *Doratodon*.

**Analysis with Doratodon_S_T OTU (Fig. S4)**

The analysis of the combined taxon based on the MTM PAL 2024.159.1 skull and the type locality material retained 70 maximum parsimony trees with a length of 1780. *Doratodon* is recovered as a paralligatorid neosuchian, and as the basalmost member of a clade including *Paralligator* spp., *Yanjisuchus longshanensis*, *Scolomastax sahlsteini*, *Turanosuchus aralensis*, *Kansajsuchus extensus* and the Dzharakuduk paralligatorid, based on an anteriorly tapering dentary symphysis (Ch. 154, state 0), an unsculpted region on the dentary below the tooth row (Ch. 155, state 1), a mediolaterally compressed and vertical dentary (Ch. 160, state 0), weakly procumbent anterior dentary alveoli (Ch. 262 state 1), and the absence of a shallow fossa at the anteromedial corner of the supratemporal fenestra (Ch. 265 state 1). The sister clade of this clade within Paralligatoridae comprises *Rugosuchus nonganensis*, *Shamosuchus djadochtaensis*, *Wannchampsus kirpachi* and the Glen Rose Form. Synapomorphies linking *Doratodon* to Paralligatoridae include a V-shaped intercondylar groove on the quadrate (Ch. 170 state 1), and a sharp ridge along the lateral side of the angular (Ch. 219, state 2). *Ogresuchus furatus* is recovered as an atoposaurid in a polytomy with *Knoetschkesuchus langenbergensis*, *Knoetschkesuchus guimarotae*, the clade formed by *Aprosuchus ghirai* and *Varanosuchus sakonnakhonensis*, and the clade formed by *Sabresuchus sympiestodon*, *Alligatorium* and *Theriosuchus* spp., based on the parallel to subparallel lateral margins of the anterior half of the suborbital bar (Ch. 278, state 0). *Batrachomimus pastosbonensis* is recovered as an indeterminate neosuchian more derived than *Isisfordia duncani*, *Stolokrosuchus lapparenti*, Atoposauridae and Paralligatoridae.

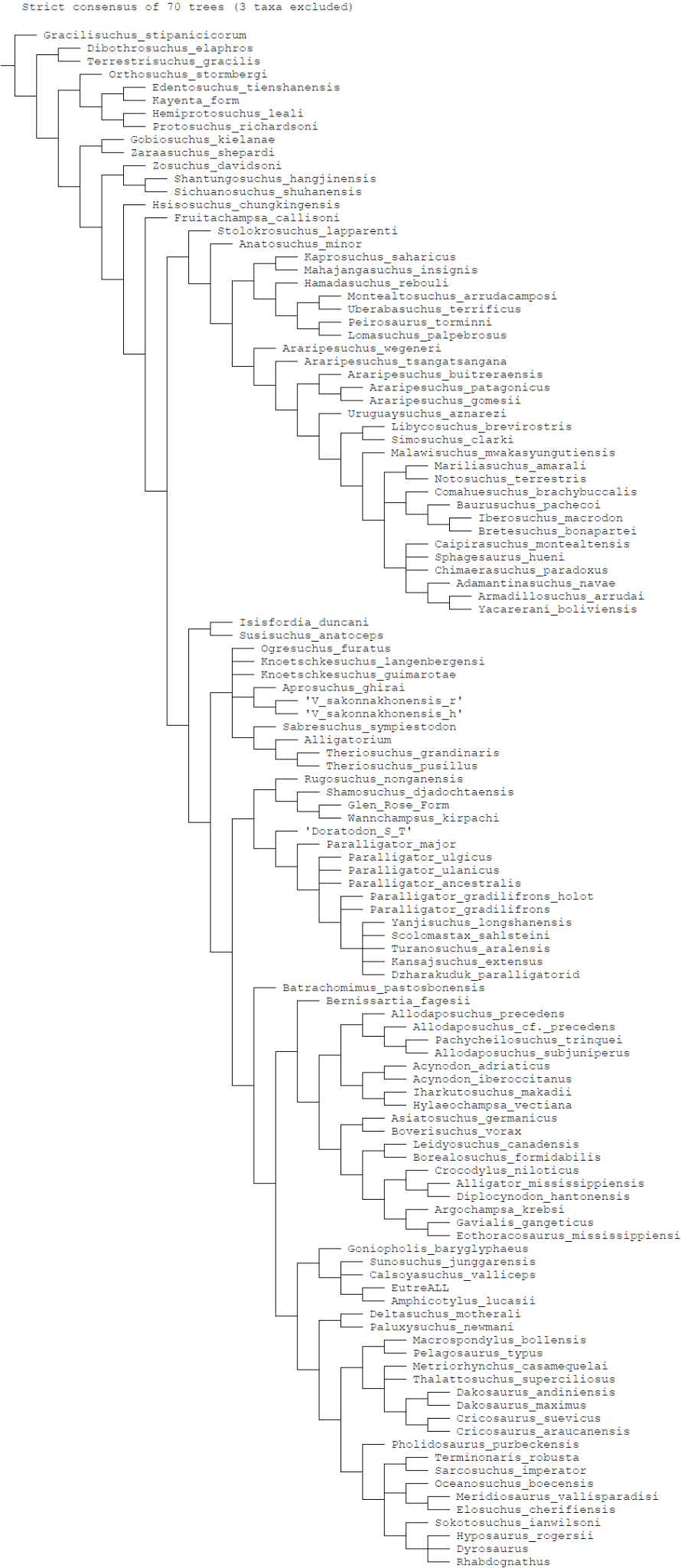

Fig. S4*:* Strict consensus of 70 MPTs from the analysis of the MTM PAL 2024.159.1 skull and Muthmannsdorf material as a single OTU in the first matrix, with *Tarsomordeo winkleri, Paralligator tersus* and *Goniopholis simus* pruned.

**Analysis with Doratodon_carcharidens_type and Doratodon_skull OTUs (Fig. S5)**

The analysis of the type locality material and the MTM PAL 2024.159.1 skull as two separate taxa retained 68 maximum parsimony trees, with a length of 1780. The two OTUs are consistently recovered as sister taxa, sharing an antorbital fenestra about half the diameter of the orbit (Ch. 67, state 1), ziphodont teeth (Ch. 120, state 1), a single laterally facing maxillary plane forming the external surface of the maxilla (Ch. 139, state 0), and a straight ventral maxillary edge (Ch. 183, state 0). *Doratodon* is consistently recovered as a paralligatorid neosuchian, and as the basalmost member of a clade including *Paralligator* spp., *Yanjisuchus longshanensis*, *Scolomastax sahlsteini*, *Turanosuchus aralensis*, *Kansajsuchus extensus* and the Dzharakuduk paralligatorid, based on an anteriorly tapering dentary symphysis (Ch. 154, state 0), an unsculpted region on the dentary below the tooth row (Ch. 155, state 1), weakly procumbent anterior dentary alveoli (Ch. 262 state 1), the absence of a shallow fossa at the anteromedial corner of the supratemporal fenestra (Ch. 265 state 1), and on some trees, a mediolaterally compressed and vertical dentary (Ch. 160, state 0). *Doratodon* shares with Paralligatoridae a V-shaped intercondylar groove on the quadrate (Ch. 170 state 1), and a sharp ridge along the lateral side of the angular (Ch. 219, state 2).

*Ogresuchus furatus* is recovered as an atoposaurid in a polytomy with *Knoetschkesuchus langenbergensis*, *Knoetschkesuchus guimarotae*, the clade formed by *Aprosuchus ghirai* and *Varanosuchus sakonnakhonensis*, and the clade formed by *Sabresuchus sympiestodon*, *Alligatorium* and *Theriosuchus* spp., based on the parallel to subparallel lateral margins of the anterior half of the suborbital bar (Ch. 278, state 0). *Batrachomimus pastosbonensis* is recovered as an indeterminate neosuchian more derived than *Isisfordia duncani*, *Stolokrosuchus lapparenti*, Atoposauridae and Paralligatoridae.

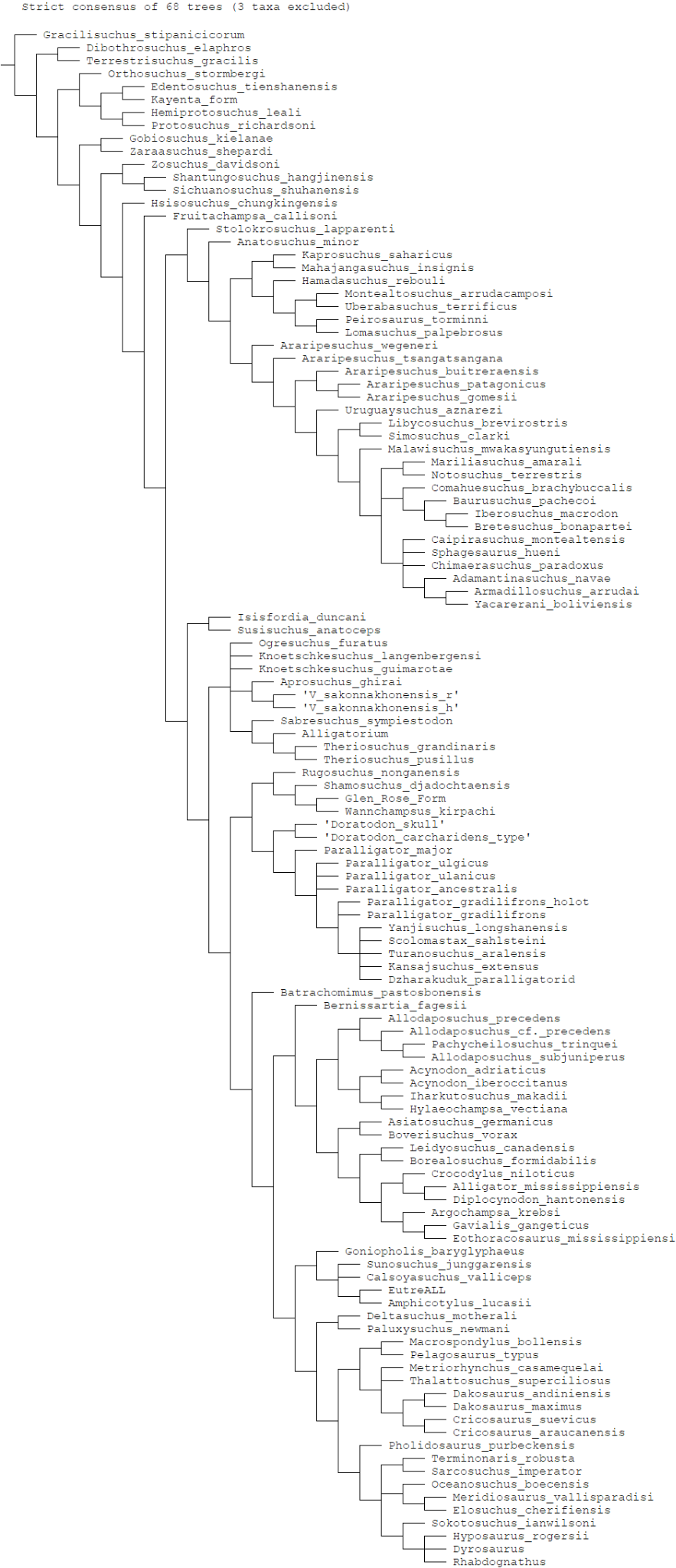

Fig. S5: Strict consensus of 68 MPTs from the analysis of the MTM PAL 2024.159.1 skull and the Muthmannsdorf material as separate OTUs in the first matrix, with *Tarsomordeo winkleri, Paralligator tersus* and *Goniopholis simus* pruned.

**Analysis with Doratodon_Ihar OTU (Fig. S6)**

The analysis of the Iharkút material retained 72 maximum parsimony trees with a length of 1778. *Doratodon* is recovered in the following positions:

A) As the sister taxon of *Sabresuchus sympiestodon* in a basal position within Mesoeucrocodylia, with which it shares a midline ridge on the dorsal surface of the frontal and parietal (Char. 22, state 1) and an unsculpted lobe on the dorsolateral process of the squamosal (Char. 35, state 1).
B) As a paralligatorid neosuchian, and the basalmost member of a clade including *Paralligator* spp., *Yanjisuchus longshanensis, Turanosuchus aralensis, Kansajsuchus extensus, Scolomastax sahlsteini* and the Dzharakuduk paralligatorid based on an anteriorly tapering dentary symphysis (Ch. 154, state 0), the presence of an unsculpted region of the dentary below the tooth row (Ch. 155, state 1), a mediolaterally compressed and vertical dentary (Ch. 160, state 0), weakly procumbent anterior dental alveoli (Ch. 262, state 1) and the lack of a fossa at the anteromedial margin of the supratemporal fenestra (Ch. 265, state 1). The clade including *Rugosuchus nonganensis, Shamosuchus djadochtaensis, Wannchampsus kirpachi* and the Glen Rose Form is sister to the aforementioned clade. *Doratodon* shares with Paralligatoridae a ventrally expanded medial quadrate condyle separated from the lateral condyle by an intercondylar groove (Ch. 170, state 1).

*Ogresuchus furatus* is consistently recovered as an indeterminate atoposaurid based on the reduced contribution of the premaxilla to the internarial bar (Ch. 4, state 1), and the parallel to subparallel anterior borders of the suborbital bar (Ch. 278, state 0).

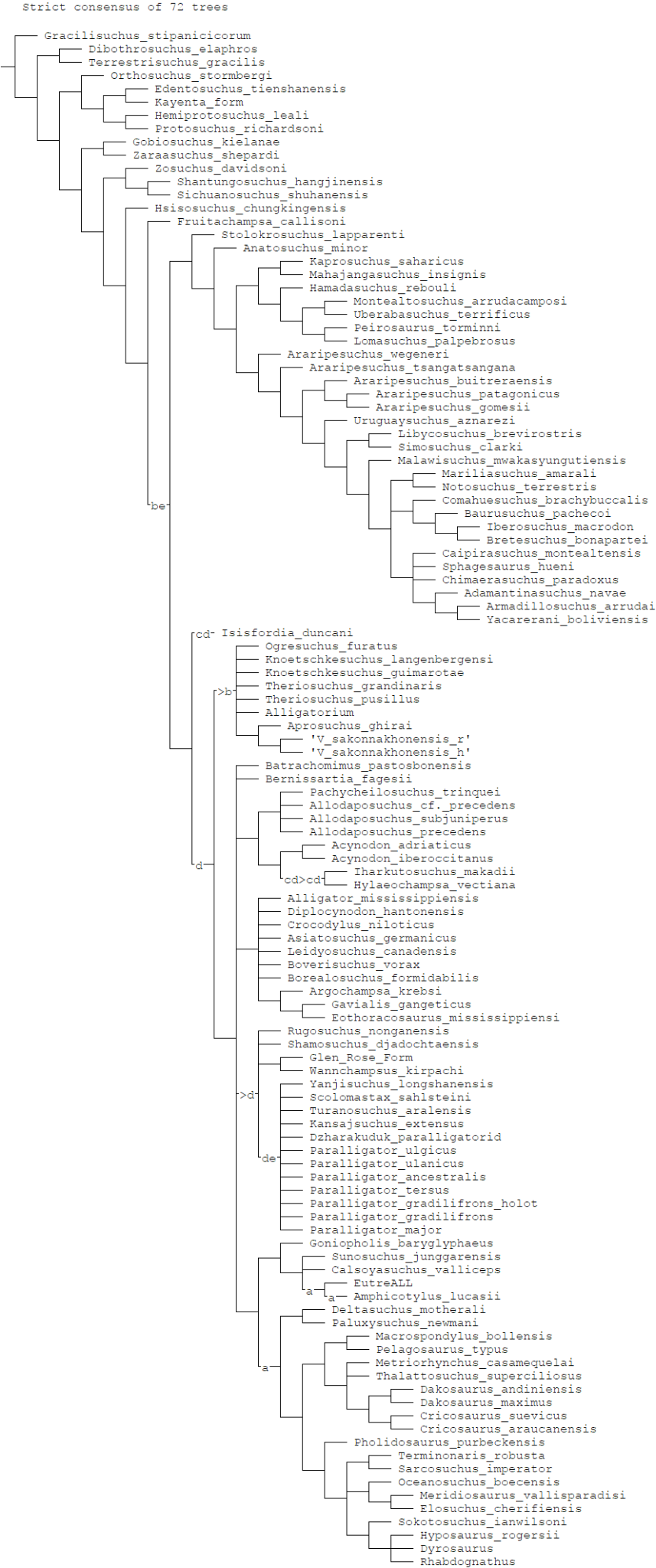

Fig. S6: Strict consensus of 98 MPTs from the analysis of the Iharkút material in the first matrix with the positions of the following taxa: a. *Goniopholis simus*, b. *Sabresuchus sympiestodon*, c. *Susisuchus anatoceps*, d. *Tarsomordeo winkleri*, e. *Doratodon*.

**Analysis with Doratodon_ALL OTU (Fig. S7)**

The analysis retained 68 maximum parsimony trees, with a length of 1780. *Doratodon* is consistently recovered as a paralligatorid neosuchian in a basal position within a clade formed by itself, *Paralligator* spp., *Yanjisuchus longshanensis*, *Turanosuchus aralensis*, *Kansajsuchus extensus*, *Scolomastax sahlsteini* and the Dzharakuduk paralligatorid, whose sister clade comprises *Rugosuchus nonganensis, Shamosuchus djadochtaensis*, the Glen Rose Form and *Wannchampsus kirpachi*. Synapomorphies linking *Doratodon* to the former clade include an anteriorly tapering dentary symphysis (Ch. 154, state 0), the presence of an unsculpted region of the dentary below the tooth row (Ch. 155, state 1), weakly procumbent anterior dental alveoli (Ch. 262, state 1), the lack of a fossa at the anteromedial margin of the supratemporal fenestra (Ch. 265, state 1), and on some trees, a mediolaterally compressed and vertical dentary (Ch. 160, state 0).

*Doratodon* shares with all of Paralligatoridae a V-shaped intercondylar groove on the quadrate (Ch. 170 state 1), and a sharp ridge along the lateral side of the angular (Ch. 219, state 2). Further synapomorphies of Paralligatoridae not preserved in *D. carcharidens* include amphicoelous or amphiplatyan cervical vertebrae (Ch. 92, state 0), the presence of appendicular osteoderms (Ch. 223, state 1), a wide and rounded olecranon process on the ulna (Ch. 260, state 1), longitudinal keels on the dorsal surface of the osteoderms restricted to the posterior edge of the osteoderm (Ch. 274, state 0), and a foramen on the premaxilla/maxilla suture near the alveolar border (Ch. 320, state 1). The clade formed by *Rugosuchus nonganensis*, *Shamosuchus djadochtaensis*, *Wannchampsus kirpachi* and the Glen Rose Form, occasionally including *Tarsomoreo winkleri*, is characterized by cranial nerves IX–XI passing through a common large foramen vagi in the otoccipital (Char. 59, state 0) and cheek teeth constricted at the base of the crown (Char. 162, state 1; also present in all other paralligatorids excluding *Doratodon carcharidens* and *Paralligator major*).

*Ogresuchus furatus* is recovered as an atoposaurid in a polytomy with *Knoetschkesuchus langenbergensis, Knoetschkesuchus guimarotae*, the clade comprising *Aprosuchus ghirai* and *Varanosuchus sakonnakhonensis*, and the clade comprising *Sabresuchus sympiestodon*, *Alligatorium* and *Theriosuchus* spp. based on the reduced contribution of the premaxilla to the internarial bar (Ch. 4, state 1), and the parallel to subparallel anterior borders of the suborbital bar (Ch. 278, state 0).

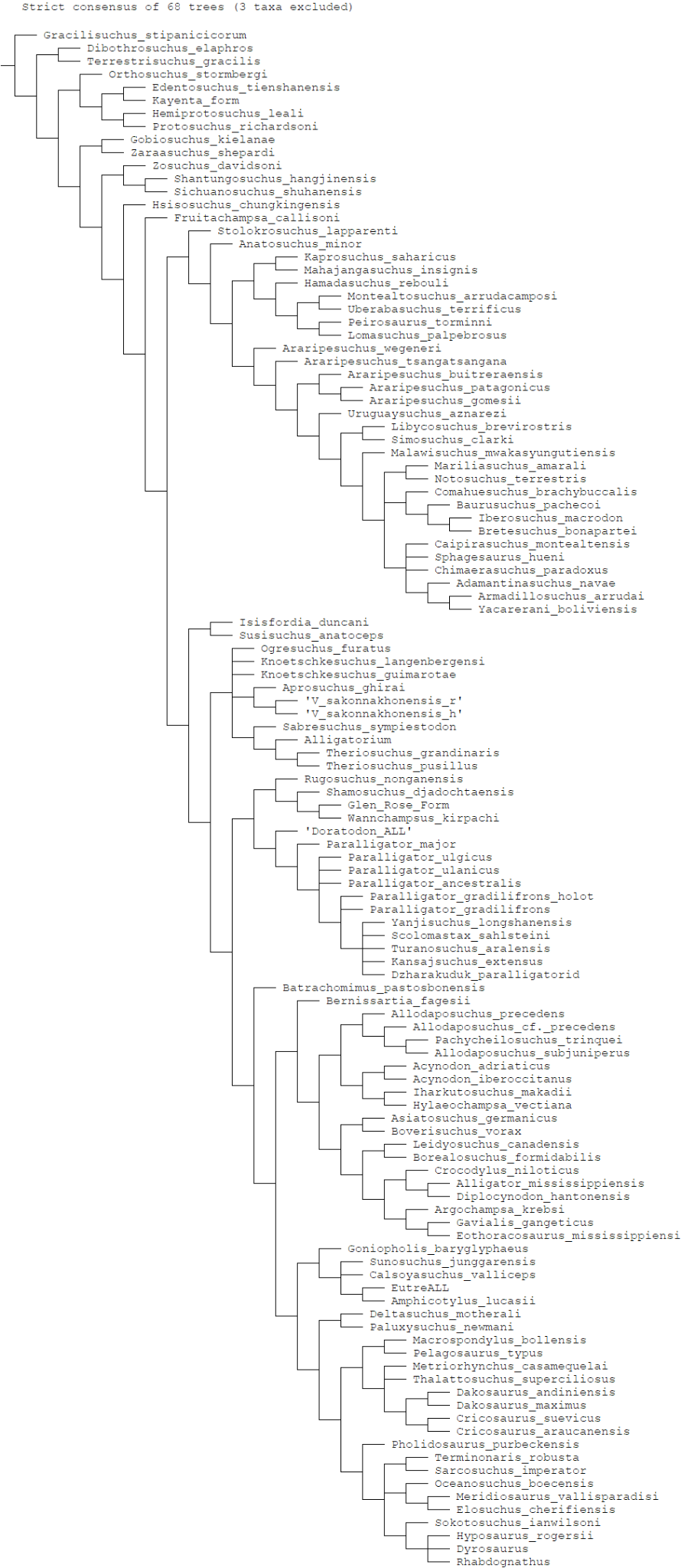

Fig. S7: Strict consensus of 68 MPTs from the analysis of the Iharkút and Muthmannsdorf material as one OTU in the first matrix, with *Tarsomordeo winkleri*, *Paralligator tersus* and *Goniopholis simus* pruned.

**Analysis with Doratodon_carcharidens_type and Doratodon_Ihar OTUs (Fig. S8)**

The analysis of the Muthmannsdorf and Iharkút material as separate OTUs recovered 62 trees with a length of 1780. The two OTUs are consistently recovered as sister taxa, sharing an antorbital fenestra about half the size of the orbit (Ch. 67, state 1), ziphodont teeth (Ch. 120, state 1), a single laterally facing maxillary plane forming the external surface of the maxilla (Ch. 139, state 0), and a straight ventral maxillary edge (Ch. 183, state 0). *Doratodon* is recovered as a paralligatorid neosuchian, and the basalmost member of the clade comprising itself, *Paralligator* spp., *Yanjisuchus longshanensis, Turanosuchus aralensis*, *Kansajsuchus extensus*, the Dzharakuduk paralligatorid and *Scolomastax sahlsteini* based on an anteriorly tapering dentary symphysis (Ch. 154, state 0), the presence of an unsculpted region of the dentary below the tooth row (Ch. 155, state 1), weakly procumbent anterior dental alveoli (Ch. 262, state 1), the lack of a fossa at the anteromedial margin of the supratemporal fenestra (Ch. 265, state 1), and on some trees, a mediolaterally compressed and vertical dentary (Ch. 160, state 0). The clade formed by *Rugosuchus nonganensis, Shamosuchus djadochtaensis,* the Glen Rose Form and *Wannchampsus kirpachi* is recovered as the sister clade of the former. *Doratodon* shares with all paralligatorids a V-shaped intercondylar groove on the quadrate (Ch. 170 state 1), and a sharp ridge along the lateral side of the angular (Ch. 219, state 2).

*Ogresuchus furatus* is recovered as an atoposaurid in a polytomy with *Knoetschkesuchus langenbergensis, Knoetschkesuchus guimarotae*, the clade comprising *Aprosuchus ghirai* and *Varanosuchus sakonnakhonensis*, and the clade comprising *Sabresuchus sympiestodon*, *Alligatorium* and *Theriosuchus* spp. based on the reduced contribution of the premaxilla to the internarial bar (Ch. 4, state 1), and the parallel to subparallel anterior borders of the suborbital bar (Ch. 278, state 0).

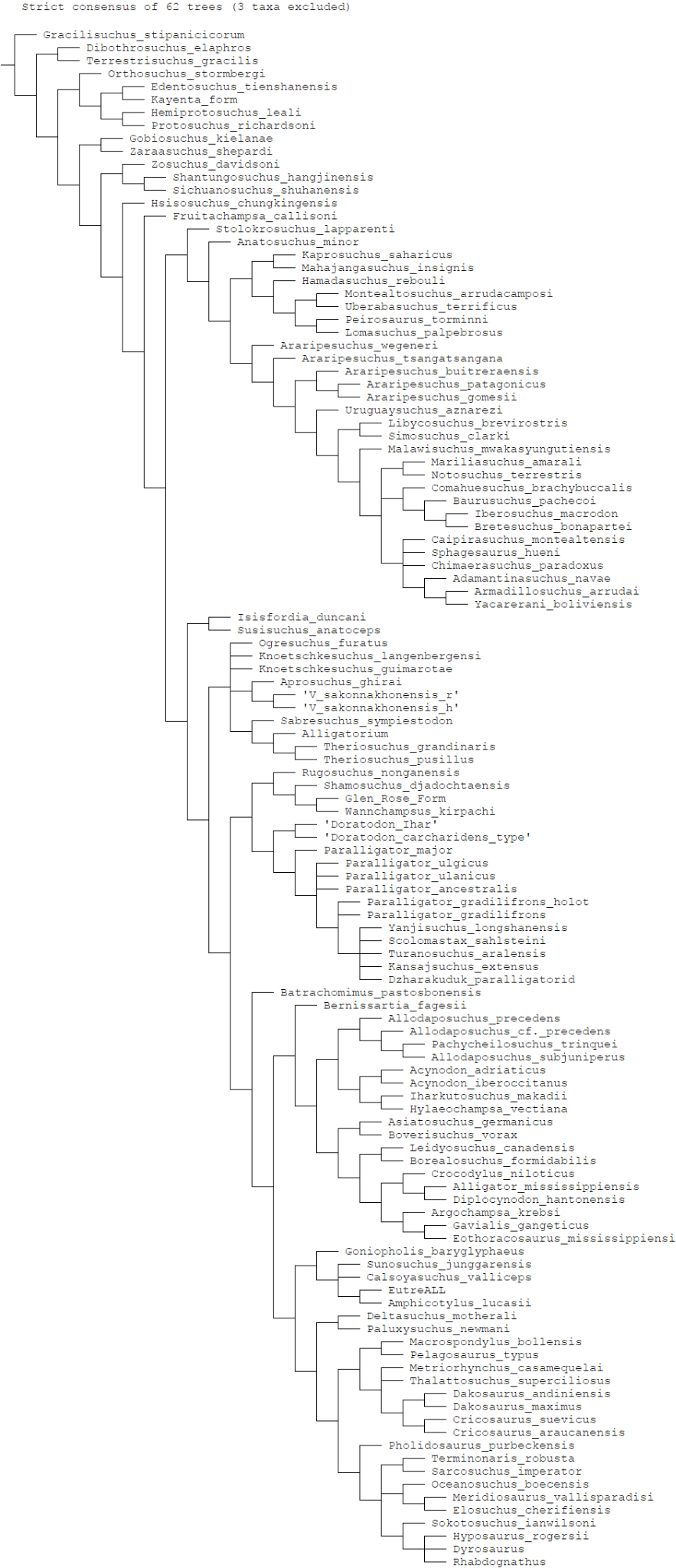

Fig. S8: Strict consensus of 62 MPTs from the analysis of the Iharkút and type locality material as separate OTUs in the first matrix with *Tarsomordeo winkleri*, *Paralligator tersus* and *Goniopholis simus* pruned.

### Dataset based on Pinheiro et al.

We used the ziphosuchian-focused dataset from Pinheiro et al.^24^, based on Pinheiro et al.^33^ and Pol et al.^34^, with the addition of the aforementioned *Doratodon* OTUs and *Ogresuchus furatus*. We set Character 127 (“Large and aligned neurovascular foramina along the lateral maxillary surface: absent (0), present (1)”) as inactive (see Dataset based on Rummy et al.). As in Pinheiro et al.^24^ and prior works, we used *Gracilisuchus stipanicicorum* as the outgroup taxon.

In our phylogenetic analyses, we used the method described in Pinheiro et al.^24^: “The memory available for the analysis was 500 MBytes, which enabled to hold 600,000 trees. Search for the minimum-length trees was conducted via New Technology using Sectorial Search, Ratchet (parameters: 25 substitutions made, or 99% swapping completed, 8 up-weighting prob., 8 down-weighting prob., and a total number of iterations of 10), Tree fusing, Driven search (50 initial addseqs., 60 times of min. length), random seed equal 0, and without collapsing trees after search. This first analysis aimed for recovering the best length number for this data matrix. Subsequently, the recovered trees were used as starting points for a Traditional Search (TBR swapping algorithm; with starting trees from RAM) and without collapsing trees after search. This second analysis aimed for recovering the maximum number of trees with the same steps recovered in the first analysis. (...) The strict consensus of the recovered [minimum-length trees] was generated. (...) All characters were treated as no additives”.

#### Analyses in the Pinheiro et al. dataset

**Analysis with Doratodon_carcharidens_type OTU (Fig. S9)**

The analysis of the *D. carcharidens* type locality material retained 600000 trees with a length of 1631. *Ogresuchus furatus* is consistently recovered as a basal notosuchian more derived than *Anatosuchus* and *Araripesuchus* spp. based on the posterior palatal interfenestral bar. *D. carcharidens* was recovered in the following positions:

A) As the sister taxon of *Theriosuchus*, forming a clade which may include *Barcinosuchus*, within Neosuchia. The clade is more basal than the clade consisting of eusuchians and paralligatorids, and may either be more basal or more derived than Goniopholidae. The two taxa share an antorbital fenestra that is less than half the diameter of the orbit, the absence of a mandibular fenestra, a ridge along the lateral angular surface, and a dentary with a single dorsal expansion posterior to which the dorsal margin is concave.
B) As the sister taxon of *Boverisuchus vorax,* in a derived position within Neosuchia. The two taxa share denticulate carinae on the teeth and a compressed and vertical dentary.
C) As the basalmost member of Sebecidae within Ziphosuchia. *D. carcharidens* shares with Sebecidae a single, laterally facing maxillary surface, and with Ziphosuchia an absent caudal intermandibular fossa and ziphodont teeth.
D) As the basalmost member of Notosuchia within Ziphosuchia. *D. carcharidens* shares with Notosuchia a dorsal dentary edge that is slightly concave or subparallel to the longitudinal axis of the skull, and with Ziphosuchia an absent caudal intermandibular fossa and ziphodont teeth.

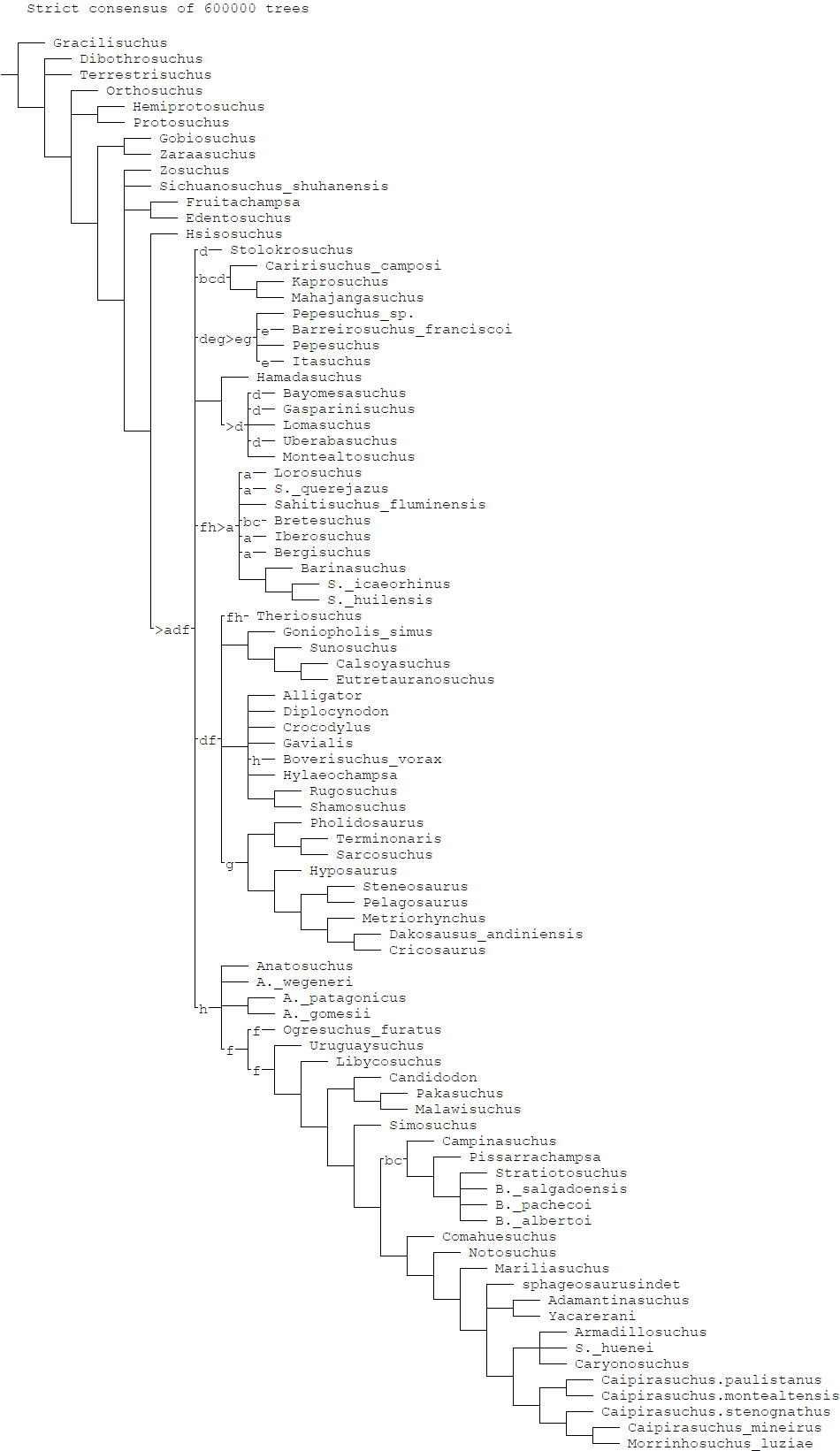

Fig. S9: Strict consensus of 600000 trees with the positions of the following wildcard-taxa: a. *Pehuenchesuchus*, b. *Ayllusuchus*, c. *Pabhwehshi*, d. *Peirosaurus*, e. *Roxochampsa*, f. *Barcinosuchus*, g. *Antaeusuchus*, e. *Doratodon carcharidens*.

**Analysis with Doratodon_skull OTU (Fig. S10)**

The second analysis of the MTM PAL 2024.159.1 *Doratodon* skull retained 152400 trees with a length of 1637. *Doratodon* is consistently recovered as the basalmost member of Neosuchia based on a lateromedially narrow supraoccipital occupying less than a third of the lateromedial width of the occipital surface, and a laterally exposed basisphenoid. *Ogresuchus furatus* is recovered as an early-diverging notosuchian more derived than *Anatosuchus* and *Araripesuchus* spp., based on a slightly constricted and posteriorly flared suborbital interfenestral bar.

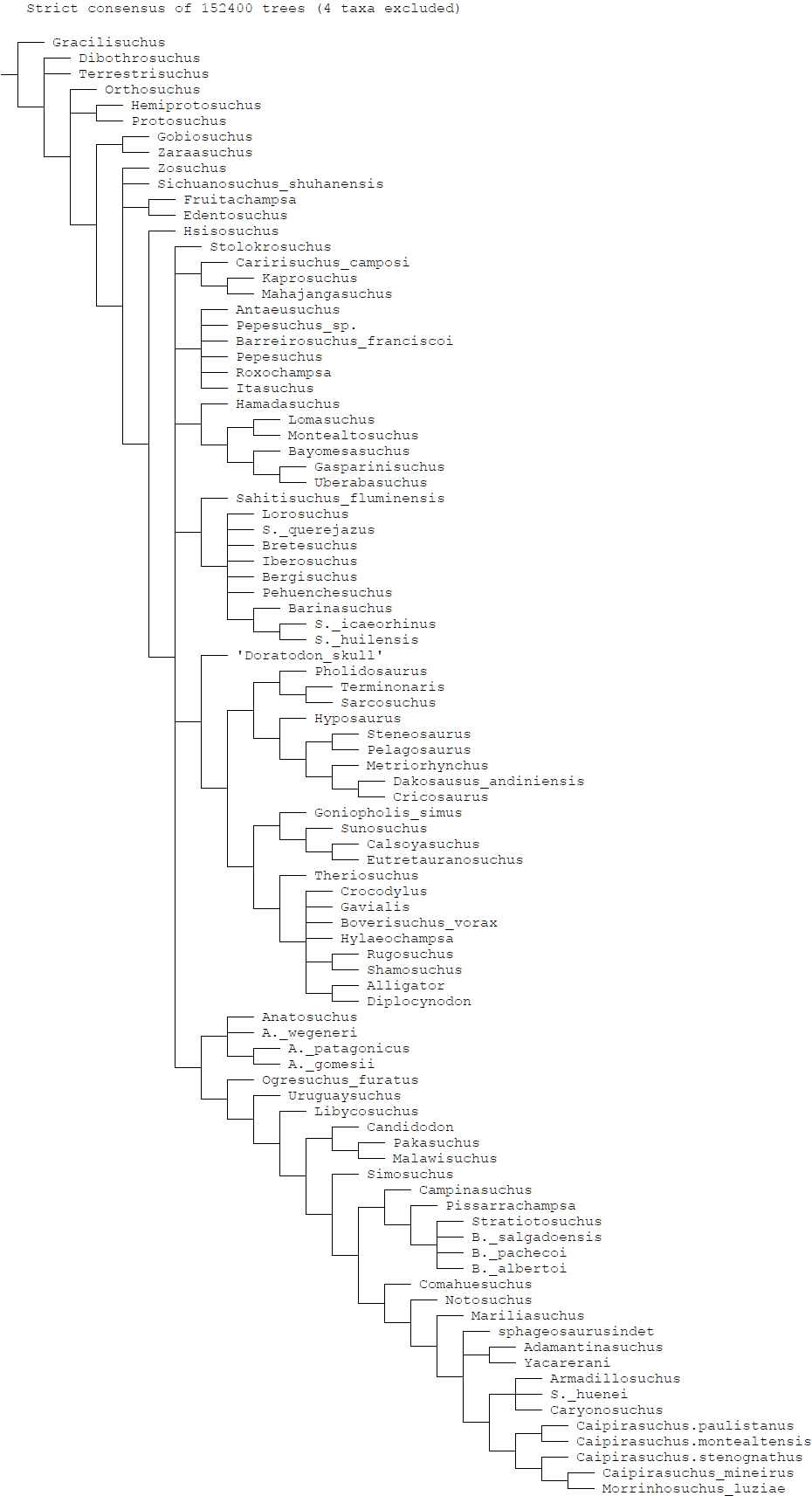

Fig. S10: Strict consensus of 152400 trees, with *Barcinosuchus, Peirosaurus, Pabhwehshi* and *Ayllusuchus* pruned.

**Analysis with Doratodon_S_T OTU (Fig. S11)**

The analysis of the MTM PAL 2024.159.1 *Doratodon* skull and the *D. carcharidens* type locality material retained 126828 trees with a length of 1542. *Doratodon* is recovered as the sister taxon of *Theriosuchus* within Neosuchia. The two form the sister clade of a derived group of neosuchians including Goniopholidae, Eusuchia and paralligatorids. *Doratodon* and *Theriosuchus* share a midline ridge on the dorsal surface of the parietal, a ventrally exposed basisphenoid, a dorsal dentary edge that expands once and is concave posteriorly, and a ridge along the lateral side of the angular, and on some trees, a present but reduced antorbital fenestra. *Doratodon* shares with derived neosuchians a dorsally transversely expanded and ventrally columnar prefrontal bar, and with all neosuchians a quadrate with two distinct faces in posterior view divided by a ridge, and a mediolaterally narrow supraoccipital. *Ogresuchus furatus* is recovered as an early-diverging notosuchian more derived than *Anatosuchus* and *Araripesuchus* spp., based on a slightly constricted and posteriorly flared suborbital interfenestral bar.

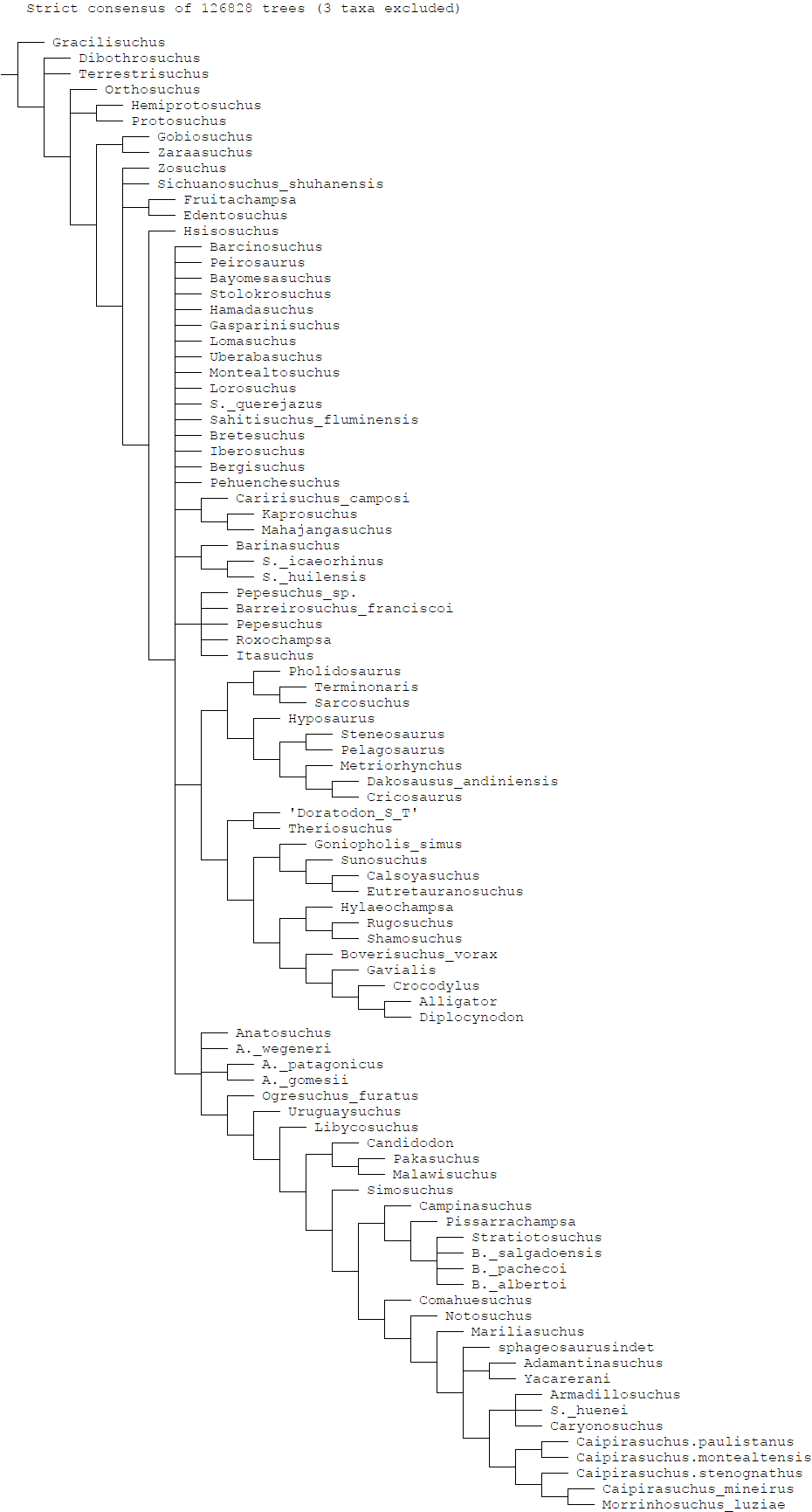

Fig. S11: Strict consensus of 152400 trees with *Antaeusuchus*, *Pabhwehshi* and *Ayllusuchus* pruned.

**Analysis with Doratodon_carcharidens_type and Doratodon_skull OTUs (Fig. S12)**

The second analysis of the MTM PAL 2024.159.1 *Doratodon* skull and the *D. carcharidens* type locality material as two separate OTUs recovered 126828 trees with a length of 1642. The two OTUs are consistently recovered as sister taxa, sharing a single laterally facing maxillary plane forming the external surface of the maxilla (Character 128, state 0). *Doratodon* as a whole is the sister taxon of *Theriosuchus* within Neosuchia, with which it forms the sister clade of a derived group of neosuchians including Goniopholidae, Eusuchia and paralligatorids. *Doratodon* and *Theriosuchus* share a midline ridge on the dorsal surface of the frontal and parietal, a ventrally exposed basisphenoid, a dorsal dentary edge that expands once and is concave posteriorly, and a ridge along the lateral side of the angular, and on some trees, a present but reduced antorbital fenestra. *Doratodon* shares with derived neosuchians a dorsally transversely expanded and ventrally columnar prefrontal bar, and with all neosuchians a quadrate with two distinct faces in posterior view divided by a ridge, and a mediolaterally narrow supraoccipital. *Ogresuchus furatus* is recovered as an early-diverging notosuchian more derived than *Anatosuchus* and *Araripesuchus* spp., based on a slightly constricted and posteriorly flared suborbital interfenestral bar and the presence of a notch on the premaxilla near the external narial border.

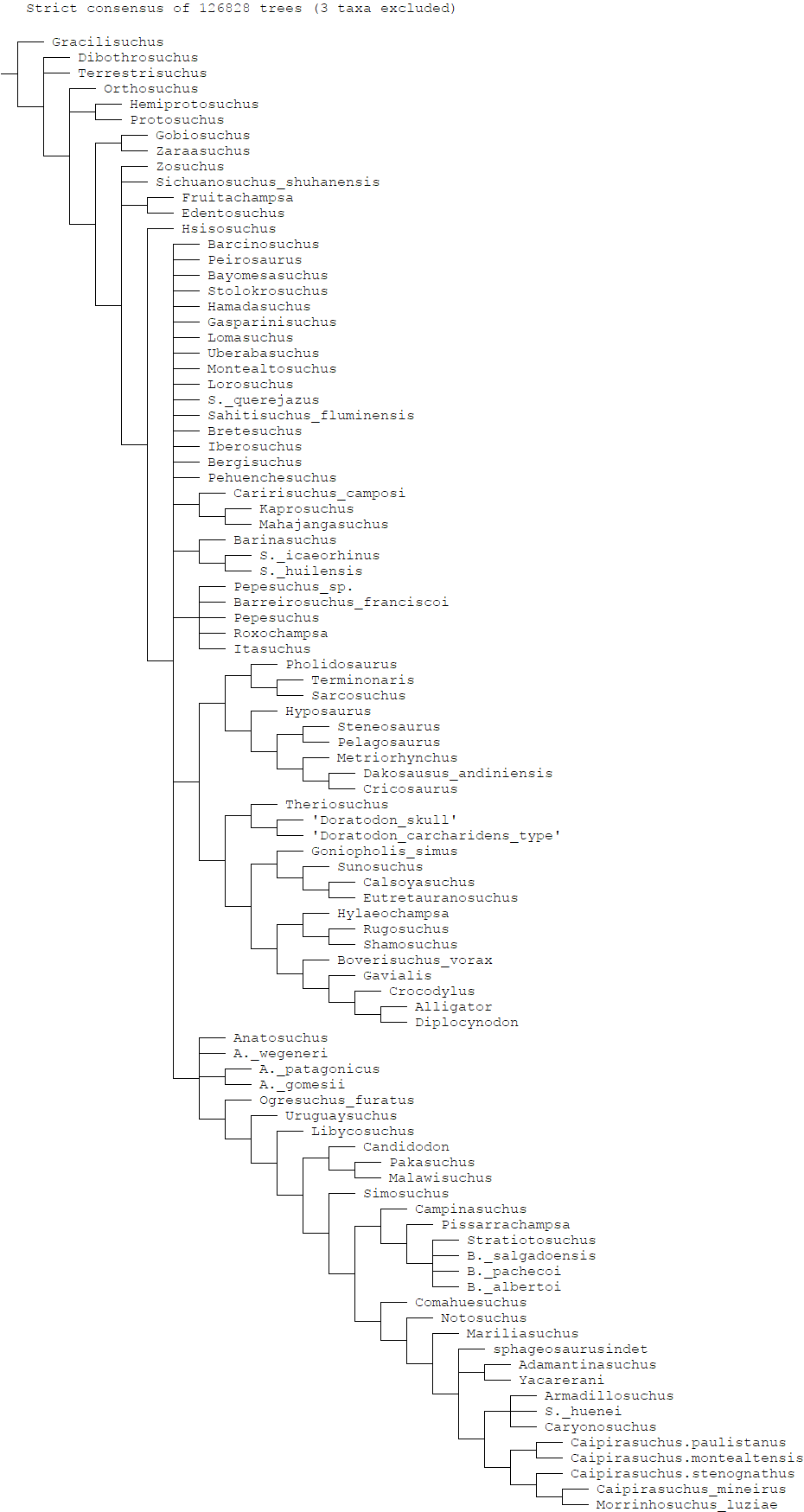

Fig. S12: Strict consensus of 126828 trees with *Antaeusuchus*, *Pabhwehshi* and *Ayllusuchus* pruned.

**Analysis with Doratodon_Ihar OTU (Fig. S13)**

The second analysis of the Iharkút material yielded 58800 trees, with a length of 1639. *Doratodon* is consistently recovered as the basalmost member of Neosuchia based on the lateral exposure of the basisphenoid on the braincase and a narrow supraoccipital less than a third of the width of the occipital surface. *Ogresuchus furatus* is recovered as an early diverging notosuchian more derived than *Araripesuchus* spp., based on the posterior flaring of the suborbital bar.

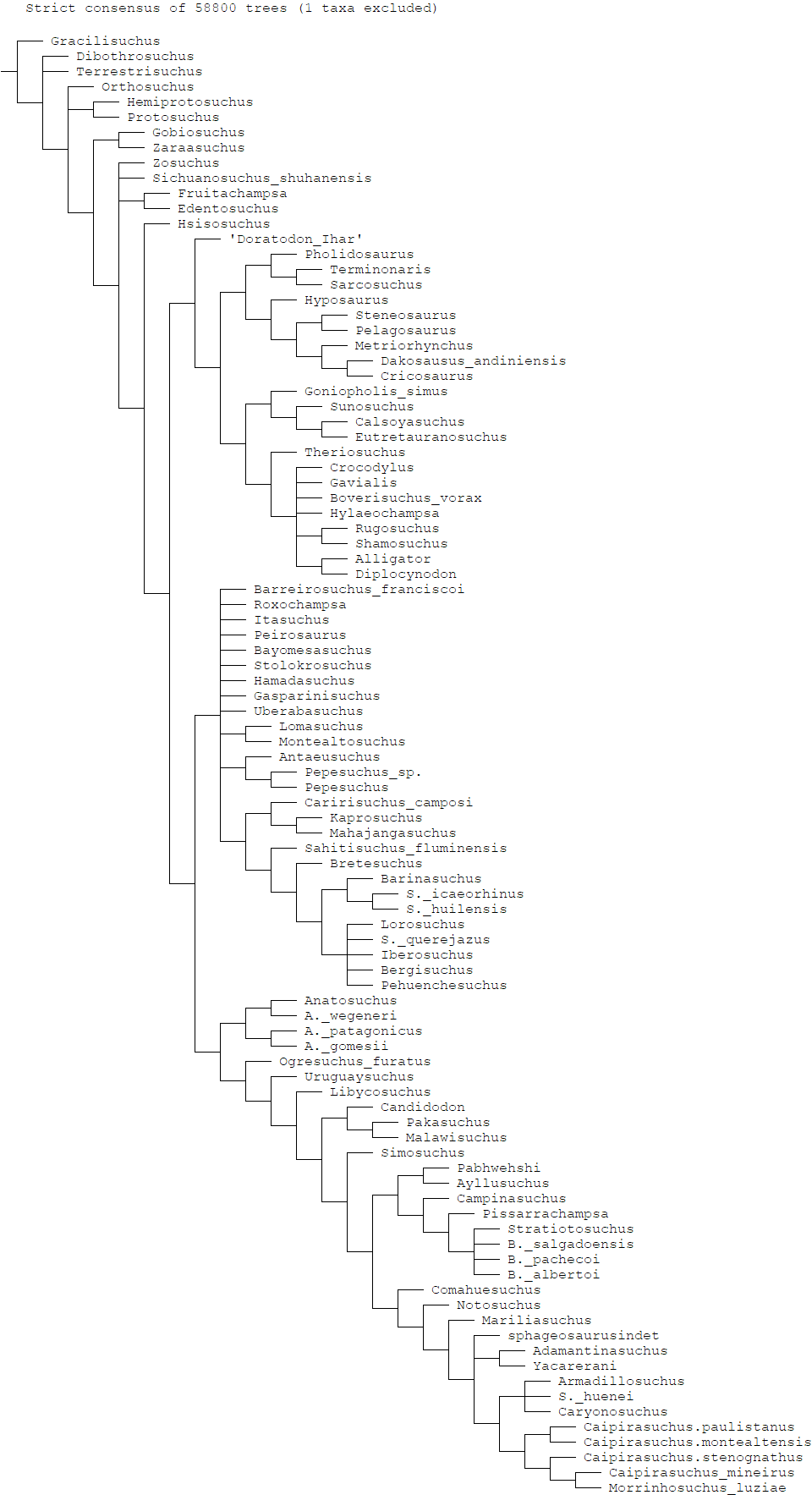

Fig. S13: Strict consensus of 58800 trees with *Barcinosuchus* pruned.

**Analysis with Doratodon_ALL OTU (Fig. S14)**

The second analysis of the type locality and Iharkút material as a combined taxon retained 126828 trees with a length of 1642. *Doratodon* is consistently recovered as the sister taxon of *Theriosuchus* within Neosuchia, with which it forms the sister clade of the clade comprising goniopholids, eusuchians and paralligatorids. *Doratodon* and *Theriosuchus* share an exposed basisphenoid on the ventral surface of the braincase, a midline ridge on the frontal and parietal, and a sharp ridge along the lateral surface of the angular. *Ogresuchus* is consistently recovered as a basal notosuchian more derived than *Araripesuchus* spp., based on a slightly constricted and posteriorly flared posterior suborbital bar and a notch on the premaxilla near the lateral edge of the external narial border.

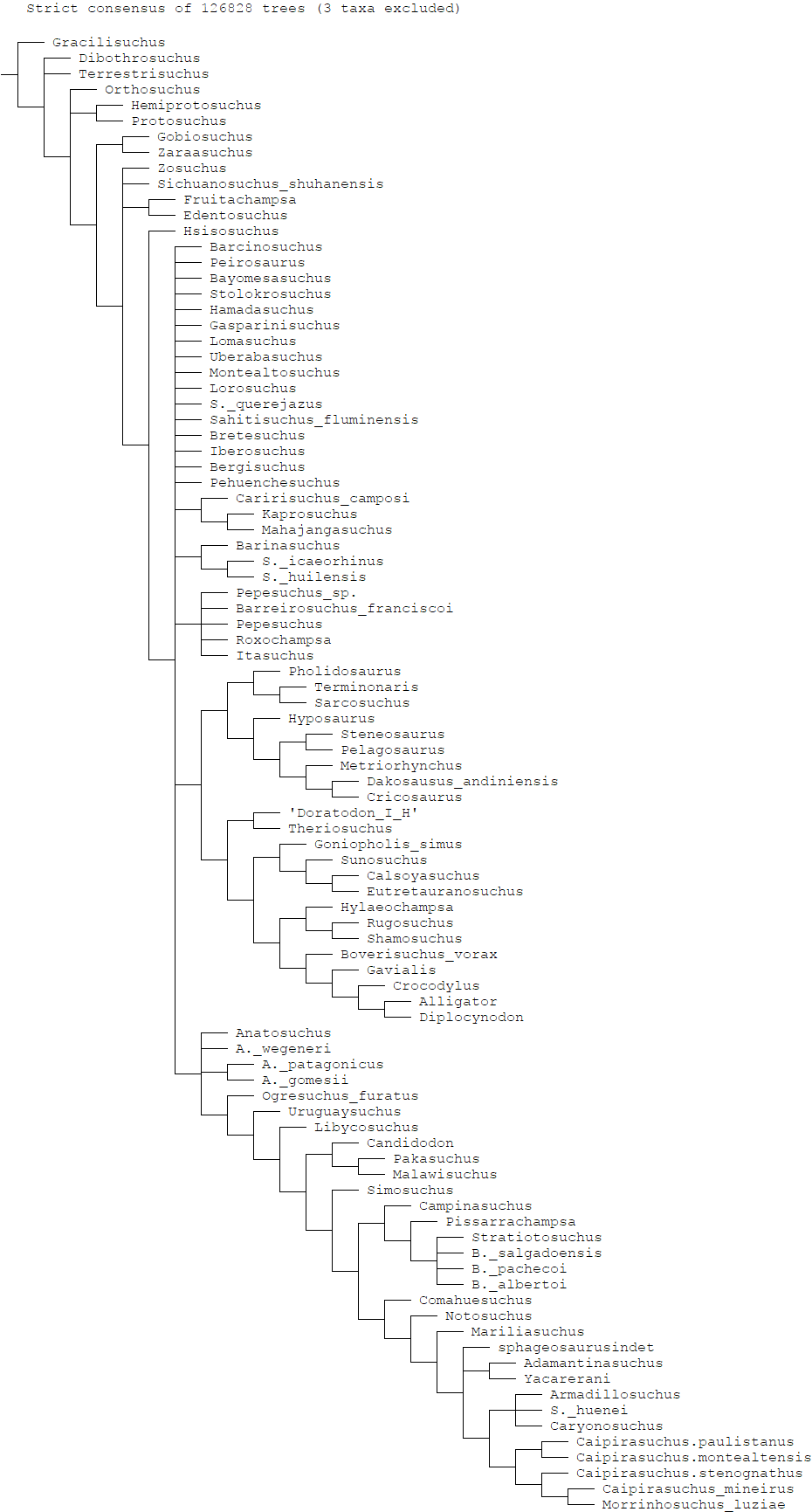

Fig. S14 Strict consensus of 126828 trees, with *Ayllusuchus*, *Pabhwehshi* and *Antaeusuchus* pruned.

**Analysis with Doratodon_carcharidens_type and Doratodon_Ihar OTU (Fig. S15)**

The analysis of the Muthmannsdorf and Iharkút material as two separate taxa yielded 119808 trees, with a length of 1640. The two *Doratodon* taxa appear as sister taxa in a clade with *Theriosuchus*, which is the sister clade of derived neosuchians including Goniopholidae, eusuchians and paralligatorids. The clade composed of *Doratodon* and *Theriosuchus* is characterized by the presence of a midline ridge on the frontal and parietal, an exposed basisphenoid on the ventral surface of the braincase, a sharp ridge along the lateral surface of the angular, and a dentary that dorsally expands once, and is concave posterior to that. The two *Doratodon* share the presence of large foramina along the alveolar edge of the dentary (Ch. 342, state 1). *Ogresuchus furatus* is recovered as a basal notosuchian more derived than *Anatosuchus* and *Araripesuchus* spp. based on a slightly constricted and posteriorly flared posterior suborbital bar.

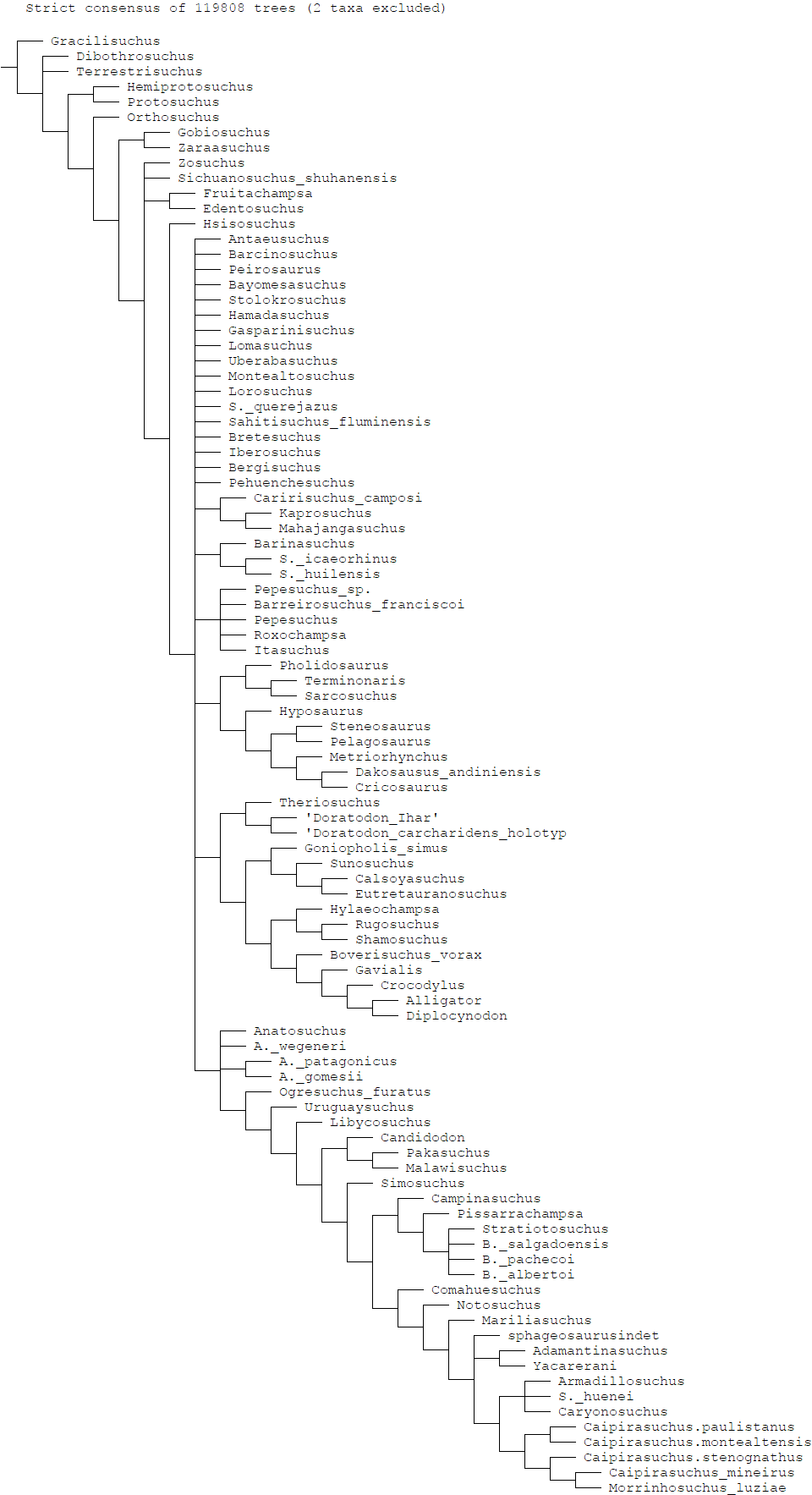

Fig. S15: Strict consensus of 119808 trees with *Ayllusuchus* and *Pabhwehshi* pruned.
